# Calprotectin protects *Staphylococcus aureus* in coculture with *Pseudomonas aeruginosa* by attenuating quorum sensing and decreasing the production of pseudomonal antimicrobials

**DOI:** 10.1101/2025.04.14.648724

**Authors:** Wei H. Lee, Amanda G. Oglesby, Elizabeth M. Nolan

## Abstract

*Pseudomonas aeruginosa* and *Staphylococcus aureus* cause debilitating polymicrobial infections in diverse patient populations. Studies of these bacterial pathogens in coculture have shown that environmental variables including Fe availability and the host-defense protein calprotectin (CP) impact coculture dynamics. To decipher how CP modulates interactions between *P. aeruginosa* and *S. aureus*, we employed dual-species RNA-seq to examine the transcriptional responses of both pathogens in coculture to CP treatment and metal depletion. Analysis of these responses revealed that, for both *P. aeruginosa* and *S. aureus*, CP treatment not only induced gene expression changes consistent with single- and multi-metal starvation responses, but also induced gene expression changes that were not observed under metal limitation. For *P. aeruginosa*, CP treatment induced gene expression changes pointing to a shift in chorismate flux away from alkylquinolone and phenazine biosynthesis towards folate biosynthesis. These observations were consistent with decreased production of alkylquinolones by *P. aeruginosa*, including the potent anti-staphylococcal alkylquinolone N-oxides. CP treatment afforded perturbed levels of two quorum sensing molecules, 3-oxo-C_12_-homoserine lactone and C_4_-homoserine lactone, produced by *P. aeruginosa*. In addition, CP treatment enhanced the ability of *S. aureus* to mount Fe starvation responses and caused *S. aureus* to express host virulence genes. This analysis illuminated physiological consequences of CP treatment that extend beyond metal starvation and that these consequences impact interspecies interactions. Our findings provide a working model in which CP effectively disarms *P. aeruginosa* by inhibiting the production of anti-staphylococcal factors and boosts the ability of *S. aureus* to protect itself from attack.

**Importance:** The innate immune protein calprotectin (CP) defends the host against bacterial pathogens by sequestering multiple essential nutrient metal ions at infection sites. In addition to this role in nutritional immunity, CP promotes the survival of *Staphylococcus aureus* in coculture with *Pseudomonas aeruginosa*, an effect that is independent of its metal-sequestering function. In this work, we sought to understand how CP modulates this interspecies interaction by evaluating the transcriptional responses of *P. aeruginosa* and *S. aureus* to CP and metal limitation in cocultures. Our study revealed that CP attenuates the ability of *P. aeruginosa* to attack *S. aureus* with anti-staphylococcal factors and enhances the capacity of *S. aureus* to withstand this assault, effects that are not recapitulated by metal limitation. This work provides new understanding of how CP modulates microbial interactions that are relevant to human health.

## Introduction

*Pseudomonas aeruginosa* and *Staphylococcus aureus* are two bacterial pathogens of clinical concern owing to their widespread prevalence, ability to colonize and thrive within the host environment and resistance against available antimicrobial therapies (1–3). *P. aeruginosa* and *S. aureus* share infection niches, including chronic wounds and the lungs of cystic fibrosis (CF) patients (4, 5). Co-infections by *P. aeruginosa* and *S. aureus* exacerbate the severity of the infection (6–8), and interactions between these two bacterial pathogens increase the tolerance of both *P. aeruginosa* and *S. aureus* to antibiotic treatment (4, 5, 9–12). However, our current understanding of how host immunity and the host environment impact interactions between these two bacterial pathogens is limited (13).

*P. aeruginosa* and *S. aureus* interactions are antagonistic, with *P. aeruginosa* outcompeting *S. aureus* by secreting various anti-staphylococcal factors (14–16). Multiple virulence factors and exoproducts contribute to the anti-staphylococcal activity of *P. aeruginosa* against *S. aureus*, including LasA, LasB and phenazines such as pyocyanin (PYO) (17). Production of anti-staphylococcal factors by *P. aeruginosa* is controlled by a hierarchy of quorum-sensing (QS) metabolites, autoinducers, and their associated regulators (18–23). Two prominent autoinducer–regulator systems involved in this hierarchical QS cascade (24–27) are the 3-oxo-C_12_-HSL/LasIR (18, 28) and the C_4_-HSL/RhlIR systems (18, 29). When activated, the 3-oxo-C_12_-HSL/LasIR system triggers the activation of downstream virulence factors and systems (30–33), including the production of alkaline protease, staphylolysin (LasA) (34, 35), and upregulation of genes encoding the C_4_-HSL/RhlR system (19). Activation of the C_4_-HSL/RhlIR system is essential for rhamnolipid biosynthesis and the production of elastase (LasB) (36–38) and PYO (39). *P. aeruginosa* also produces multiple alkylquinolones (AQs) which contribute to the anti-staphyloccocal activity of *P. aeruginosa* both as QS molecules and as direct anti-staphylococcal metabolites (40–44). The combination of multiple molecular factors drives *S. aureus* towards fermentative metabolism and reduces *S. aureus* viability (11, 15, 45, 46).

The host environment undoubtedly affects interactions between *P. aeruginosa* and *S. aureus*. One environmental variable is nutrient availability. *P. aeruginosa* and *S. aureus* have a metabolic Fe requirement, and the host lowers Fe availability to starve invading pathogen in a process termed nutritional immunity (47, 48). In response to host-imposed Fe limitation, *P. aeruginosa* and *S. aureus* express dedicated transporters (49, 50), siderophores (51–54), and heme uptake machinery (55, 56) to compete for this nutrient. Prior studies of *P. aeruginosa* and *S. aureus* cocultures have examined the importance of metal availability, revealing that Fe starvation enhances anti-staphylococcal activity of *P. aeruginosa* toward *S. aureus* (57, 58). The host protein calprotectin (CP) contributes to nutritional immunity by sequestering multiple divalent transition metal ions including Fe(II) and elicits single- and multi-metal starvation responses in these two bacterial pathogens (59–65). CP was also shown to promote *S. aureus* survival in coculture with *P. aeruginosa*, which was attributed to reduced production of anti-staphylococcal factors resulting from CP-mediated metal limitation (66). Combined, these studies revealed an apparent dichotomy in the field, wherein CP sequesters Fe(II) and induces Fe-starvation responses in both pathogens, while also providing a protective effect on *S. aureus* when cocultured with *P. aeruginosa* (66–68). Our recent work demonstrated that the protective effect of CP on *S. aureus* is metal-independent, indicating that the CP protein scaffold directly impacts coculture dynamics (68). Our findings also suggested that perturbed production of PYO and the siderophore pyochelin (PCH) by *P. aeruginosa* in the presence of CP may arise due to additional effects of CP that extend beyond its metal sequestering ability (68). Others have shown that CP interacts physically with *P. aeruginosa* and *S. aureus* during coculture (69), though the impact of these interactions on the anti-staphylococcal activity of *P. aeruginosa* is unknown. Taken together, these studies illustrate the complex and multifactorial effects of the CP protein scaffold and metal limitation on coculture outcomes, necessitating a new model for how CP and metal availability modulate interspecies dynamics between *P. aeruginosa* and *S. aureus*.

To support the development of such a model, we utilized dual-species RNA-seq to evaluate the global transcriptional responses of *P. aeruginosa* and *S. aureus* in coculture to CP treatment and metal depletion. We report that CP treatment elicits transcriptional responses in both bacterial pathogens that shape coculture outcomes and that were not observed in metal-depleted cocultures. In *P. aeruginosa*, CP treatment induced gene expression changes indicating redirected chorismate flux, perturbed levels of QS effectors, and decreased production of AQs, effectively reducing the anti-staphylococcal activity of *P. aeruginosa*. Consistent with these observations, the presence of CP led to decreased gene expression responses related to membrane damage and cell stress while enhancing Fe starvation responses by *S. aureus*. CP treatment also increased the expression of *S. aureus* genes associated with host virulence. Our findings support a model in which *P. aeruginosa* functions as an attacker that antagonizes *S. aureus*, the defender, occurring primarily through the action of alkylquinolone N-oxides. By perturbing QS production and decreasing AQ production, CP effectively disarms *P. aeruginosa* and promotes the survival of *S. aureus* in coculture. Collectively, our results show that the CP protein scaffold significantly impacts coculture dynamics between these two pathogens, demonstrating that components of host immunity may impact pathogen–pathogen interactions in ways outside their known function.

## Results and Discussion

### Experimental considerations for dual-species RNA-seq

The study design leveraged insights from prior investigations of the effects of CP treatment and metal depletion on the growth dynamics of *P. aeruginosa* and *S. aureus* cocultures, utilizing the same coculture conditions as previously described (68). Briefly, *P. aeruginosa* strain UCBPP-PA14 (hereafter PA14) and *S. aureus* strain USA300 JE2 (hereafter JE2) were grown in a chemically defined medium (CDM) prepared from trace metals basis reagents used in prior studies of both species in monocultures or cocultures (59–61, 68). This base medium is used to prepare media with defined metal concentrations. Metal-replete CDM is supplemented with 0.3 μM Mn, 5 μM Fe, 0.1 μM Cu, 0.1 μM Ni, and 6 μM Zn. These metal concentrations are representative of physiologically relevant metal levels in sputum samples from CF patients (70, 71). The omission of one or more metals (Mn, Fe, Zn, or all three metals) from this mixture allows for comparisons between the effects of CP treatment and either single- or multi-metal depletion on *P. aeruginosa* and *S. aureus* in monoculture and coculture. We performed dual-species RNA-seq on cocultures of these two bacterial pathogens grown in metal-replete, Mn-depleted, Fe-depleted, Zn-depleted and metal-depleted CDM (depleted of Mn, Fe and Zn), and metal-replete CDM supplemented with a physiologically relevant concentration of CP (20 μM) (68, 72, 73). We compared the transcriptional responses of the cocultures exposed to metal depletion with those observed for CP treatment. We also performed RNA-seq on *P. aeruginosa* and *S. aureus* monocultures treated with CP (20 μM) to identify transcriptional responses occurring specifically in coculture or in monoculture of either species as well as responses common to both culture types.

We identified the 6–8 h period as a potential window for sample collection based on our prior coculture growth and metabolite time-course studies. In this timeframe, Fe starvation responses including appreciable production of the pseudomonal siderophores pyoverdine and PCH and decreased production of phenazines occurred from 6 h onwards in cultures treated with CP (68). To determine the appropriate timepoint for RNA-seq sample collection, real-time PCR was used to confirm sample reproducibility by validating consistent transcript levels for both *P. aeruginosa* and *S. aureus* cocultured in Fe-depleted medium and metal-replete medium treated with or without CP (68). Across the conditions tested, *P. aeruginosa* RNA abundance remained high as judged from transcript levels of the housekeeping gene, 16S. However, the reproducibility of *S. aureus* RNA transcripts from cocultures grown in the absence of CP declined significantly past the 6 h timepoint as judged from highly variable and often trace levels of the *S. aureus* housekeeping gene *sigA* (**Table S1**), indicative of significant RNA degradation as *P. aeruginosa* anti-staphylococcal activity proceeded. Consequently, the 6 h timepoint was selected for RNA-seq. For differential expression analysis, metal-replete CDM functioned as the (untreated) control. The proportions of differentially expressed (DE) genes for *P. aeruginosa* and *S. aureus* are presented in **Table S2**. Key statistics of the RNA-seq dataset, including the sequenced read count and the number of unique features sequenced (74), are presented in **Table S3**. Herein, we summarize multi-metal starvation responses induced by CP in cocultures of *P. aeruginosa* and *S. aureus*, and describe how CP modulates interspecies dynamics between both bacterial pathogens. Additional transcriptional responses of *P. aeruginosa* unique to each experimental condition are presented in the accompanying **Supplementary Discussion**. The complete list of DE *P. aeruginosa* and *S. aureus* genes are presented in the accompanying **Supplementary Files**.

### CP elicits multi-metal starvation responses in *P. aeruginosa* cocultured with *S. aureus*

We examined the top 600 DE genes across all culture conditions with Venn analysis to compare responses resulting from CP treatment and metal depletion. For *P. aeruginosa* genes DE in response to CP treatment and Fe depletion, about 48% of upregulated genes (**Figure 1A**) and 40% of downregulated genes (**Figure 1B**) were common to both conditions, demonstrating significant overlap. For genes DE in response to CP treatment and Zn depletion, 47% of upregulated genes (**Figure 1A**) and all downregulated genes (**Figure 1B**) were common to both conditions. For cocultures grown in Mn-depleted CDM, only 42 *P. aeruginosa* genes fell within the top 600 DE genes across all culture conditions (**Figures S1A** and **S1B**), and nearly all genes observed to be upregulated in response to Mn-depletion were also upregulated in CP-treated cocultures (**Figure S1A**). Transcriptional responses common to CP treatment and the depletion of Fe, Zn or Mn were also recapitulated in metal-depleted cocultures (**Figures S1 – S3**). While most transcriptional responses (∼87%) of *P. aeruginosa* in monoculture to CP treatment overlapped with the transcriptional responses for *P. aeruginosa* in coculture to CP treatment, approximately 9% of upregulated genes and 16% of downregulated genes were found to be unique to either monoculture or coculture (**Figures S4A** and **S4B**).

**Figure 1.**
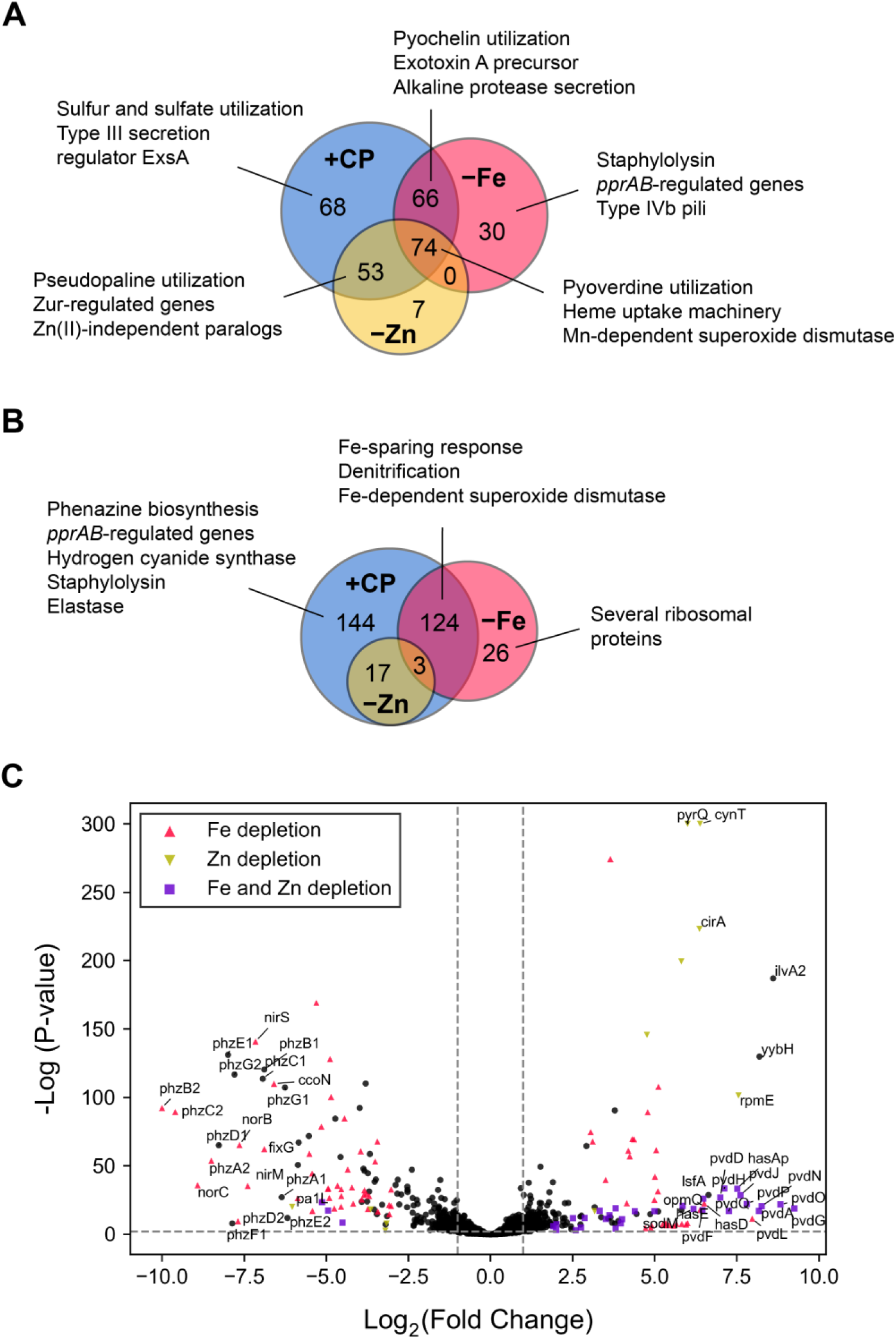
CP induces multi-metal starvation responses by *P. aeruginosa* cocultured with *S. aureus*. Venn analyses reveal significant overlap of upregulated (**A**) and downregulated (**B**) *P. aeruginosa* genes in cocultures treated with CP and cocultures grown in Fe-depleted or Zn-depleted CDM. The top 600 DE genes across all culture conditions were used for Venn analyses. (**C**) Volcano plot of DE changes in response to CP treatment. Genes with similar DE patterns in response to Fe depletion, Zn depletion or both Fe and Zn depletion are denoted as colored shapes. A threshold cutoff Log_2_(Fold Change) of 1 was employed.

**Figure 2.**
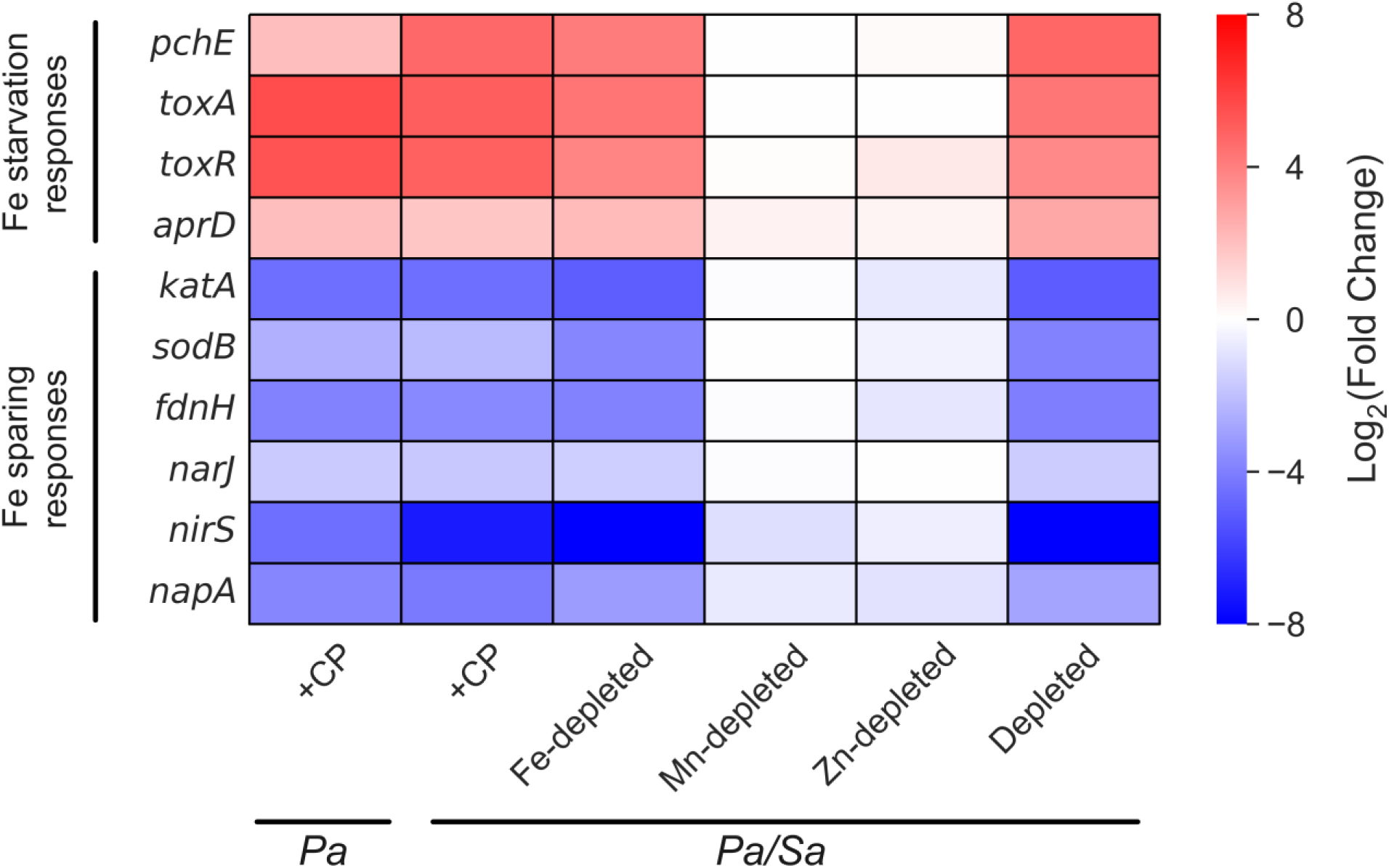
CP elicits Fe starvation responses and Fe sparing responses by *P. aeruginosa* cocultured with *S. aureus*. DE heatmap of *P. aeruginosa* genes associated with Fe starvation responses and Fe sparing responses. *Pa* indicates *P. aeruginosa* monoculture and *Pa/Sa* indicates the coculture.

To further probe similarities and differences between CP treatment and metal depletion, we performed functional enrichment analyses. Overrepresentation analysis of DE genes revealed considerable overlap between the effects of CP treatment and metal depletion for *P. aeruginosa* in coculture (**Figures S5A** and **S5C**), whereas genes from several distinct clusters of Gene Ontology (GO) terms responded uniquely to CP treatment or the depletion of Fe, Mn or Zn (**Figures S5B** and **S5D**). Genes encoding for the Mn-dependent superoxide dismutase SodM (75), the heme acquisition protein HasAp (76), Phu heme uptake machinery (55), and pyoverdine biosynthesis and uptake machinery (*pvd* operon) (52, 68) were upregulated in all four conditions (**Table SF1A**). No systems were downregulated in all four conditions (**Table SF1B**). Collectively, our findings demonstrate that CP treatment elicits multi-metal starvation responses in *P. aeruginosa* co-cultured with *S. aureus*, and these responses overlap considerably, but not fully, with the effects of metal depletion.

### CP induces Fe starvation responses in *P. aeruginosa* cocultured with *S. aureus*

In agreement with prior studies reporting that CP elicits transcriptional changes associated with Fe-starvation responses in *P. aeruginosa* during coculture with *S. aureus* (68) and in monoculture (59, 61, 66), CP treatment and growth in Fe-depleted CDM resulted in the upregulation of pyoverdine (*pvd*) and pyochelin (*pch*) biosynthetic machinery (53), the exotoxin A precursor *toxA* (66, 77), the transcriptional regulator *toxR* (78) and alkaline protease secretion machinery (79) (**Figures 1A**, **2** and **Table SF2A**). These observations are also consistent with previous studies of Fe starvation responses in *P. aeruginosa* (60, 80). Genes encoding for the MexEF-OprN efflux pump, which secretes the QS metabolite HHQ (81, 82) and contributes to antimicrobial resistance (83), and the Fe-independent paralog of fumarase *fumC1* (84) were also found to be upregulated in response to CP treatment and Fe depletion (**Table SF2A**).

Under conditions of Fe limitation, *P. aeruginosa* decreases the expression of Fe-containing proteins via an Fe sparing response that is mediated by PrrF small RNAs (sRNAs) (85–87). CP treatment and Fe depletion resulted in strong downregulation of the Fur-regulated catalase *katA* (88, 89), the Fe- cofactored superoxide dismutase *sodB* (90), the nitrate-inducible formate dehydrogenase *fdnIHG* (91, 92) and genes involved in denitrification (*nar*, *nir*, *nap*), indicative of Fe sparing responses under these treatment conditions (93) (**Figures 1B**, **2** and **Table SF2B**). Furthermore, CP treatment and Fe depletion decreased the expression of the cbb3-type cytochrome c oxidase *cco* (66, 94–96) and the NADH dehydrogenase *nuo* (97) (**Figure S6** and **Table SF2B**), as part of an Fe starvation response (80). CP treatment and Fe depletion also decreased the expression of the bacterial ferritin *ftnA* (98) and phenazine biosynthetic machinery (*vide infra*) (**Figure S7**).

### CP induces Zn starvation responses for *P. aeruginosa* cocultured with *S. aureus*

Consistent with Zn starvation responses (60), CP treatment and Zn depletion resulted in the upregulation of genes associated with Zn uptake machinery, including pseudopaline biosynthesis and transport (*cnt*) (99, 100), the *znu* operon (101, 102), and the Zn uptake regulator *zur* (101) (**Figures 1A**, **S8** and **Table SF3A**). Furthermore, expression of Zn(II)-independent paralogs of *rpmE* and *rpmJ* (*PA14_17700* – *PA14_17710*) (60) and a predicted Zn(II) uptake cluster (*PA14_26390* – *PA14_26420*) (102) were upregulated upon CP treatment and Zn depletion (**Figures 1A** and **S8**). In addition, a Zur-regulated cluster of Zn-independent paralogs (*PA14_72980* – *PA14_73070*) (60, 102) was among the most strongly upregulated hits in CP-treated and Zn-depleted cocultures (**Table SF3A**). This cluster contained the cell-wall amidase *amiA* (103, 104), a putative carbonic anhydrase *cynT* (60) and the Zn-independent transcription factor *dksA2* (105, 106) (**Figure S8**). Genes downregulated by CP treatment and Zn depletion included several that are known to be regulated by the *pprAB* two-component system (107) (*vide infra*) (**Table SF3B**). Unexpectedly, we found that expression of the ferrous iron uptake system *feoAB* (49, 60, 108) and the catecholate siderophore receptor *cirA* (109, 110) were upregulated in CP-treated and Zn-depleted conditions but not in Fe-depleted conditions (60) (**Figure S8**).

### CP does not induce Mn starvation responses in *P. aeruginosa* cocultured with *S. aureus*

We detected no DE of genes encoding for the putative Mn uptake proteins MntH1 and MntH2 or genes encoding for proteins known to be Mn-cofactored in CP-treated cocultures (60) (**Table SF4**). These findings are consistent with prior studies reporting that CP treatment had negligble effect on cell-associated Mn levels in *P. aeruginosa* PAO1 grown under aerobic conditions (61).

### CP induces gene expression changes associaxted with cell envelope modifications for *P. aeruginosa* cocultured with *S. aureus*

Having characterized the aforementioned metal starvation responses of *P. aeruginosa* to CP treatment, we looked to functional categories of transcriptional changes that could not be fully accounted for by metal depletion. The presence of CP resulted in DE of multiple systems associated with modifications to the cell envelope of *P. aeruginosa* in monoculture and in coculture with *S. aureus*. These gene expression changes were mostly absent for cocultures grown in Fe-depleted conditions. In agreement with prior work (68), CP upregulated the expression of genes encoding for the spermidine synthase SpeE2 (111, 112), the 4-amino-4-deoxy-L-arabinose lipid A transferase ArnT (113, 114) and the holin CidA (*PA14_19680*), although the change in expression of *cidA* fell below the DE threshold (**Figure 3**). CP treatment resulted in downregulation of the *pprAB* two-component system (**Figure 3**), which regulates a hyper-biofilm phenotype (Pel- and Psl-independent) in *P. aeruginosa* (107). Furthermore, genes known to be positively regulated by *pprAB* (107), including those encoding for structural components and assembly of the type IVb pili (115, 116), CupE fimbriae (117), and the BapA adhesin (107, 118), were downregulated by CP treatment (**Figure 3**). We observed that downregulation of genes associated with the *pprAB* regulon was partially attributable to Zn depletion. CP treatment also upregulated the expression of the *arn* operon and the *PA14_63110* – *PA14_63160* locus containing the two-component sensor/regulator *pmrAB* (119, 120), transcriptional responses which were common to Zn-depleted cocultures (**Table SF3A**). The *pprAB* two-component system has also been reported to regulate PQS biosynthetic and transport machinery (*pqsCDE*) (*vide infra*) and anthranilate synthase (*phnAB*) (107) (**Figure 3**).

**Figure 3.**
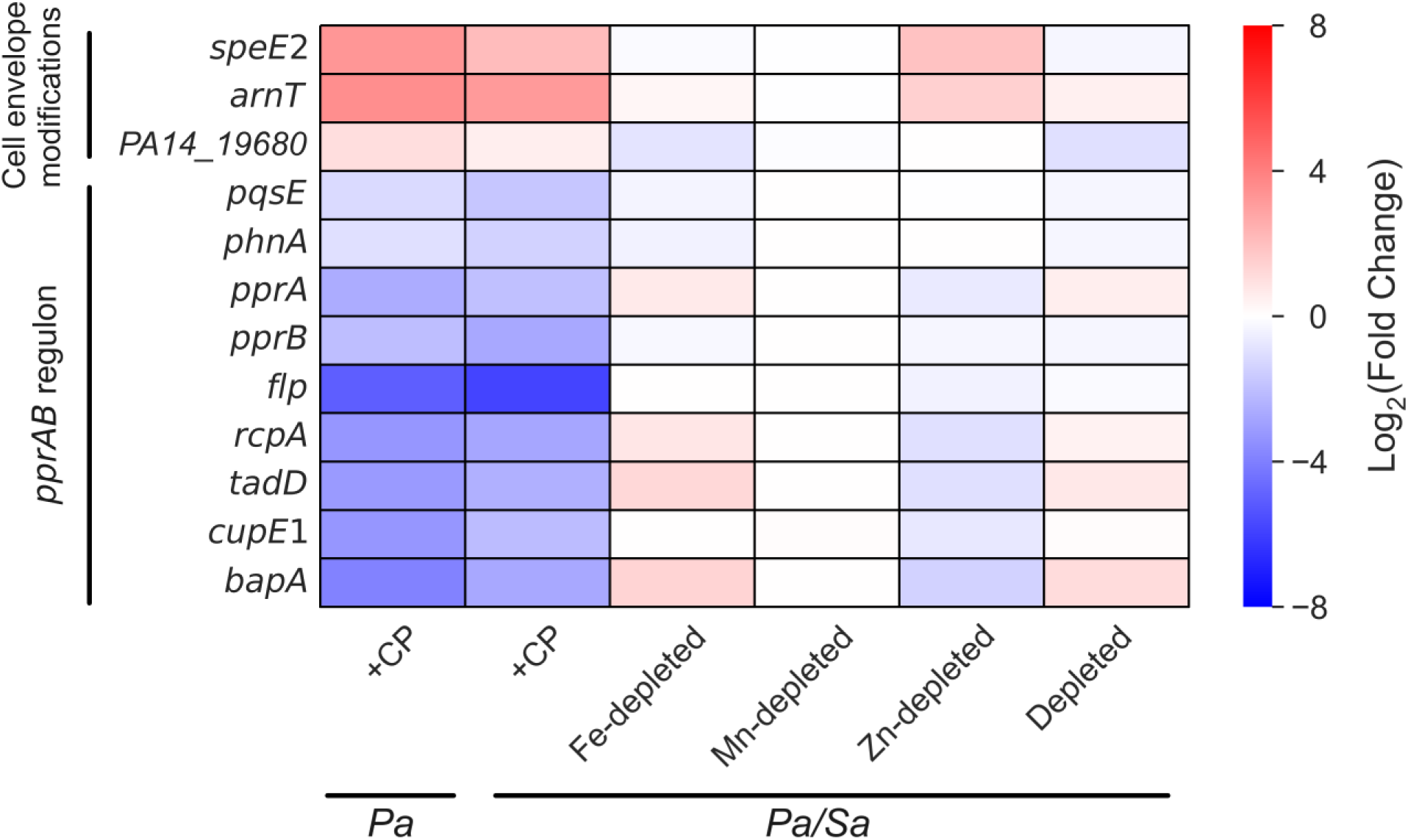
CP elicits transcriptional responses associated with cell envelope modifications for *P. aeruginosa* in coculture with *S. aureus*. DE heatmap of *P. aeruginosa* genes associated with cell envelope modifications and the *pprAB* regulon. *Pa* indicates *P. aeruginosa* monoculture and *Pa/Sa* indicates the coculture.

We also found that CP elicited transcriptional changes that were not attributable to metal depletion. CP treatment caused mild upregulation of the *cprRS* (121) and *parRS* (122) two-component systems (**Figure S9**), which are associated with *P. aeruginosa* responses to cationic peptides and antibacterials including aminoglycosides and polymyxins. Additional indications of altered membrane character/integrity in response to CP treatment included upregulation of the genes encoding for the *N*- succinyl-L-diaminopimelic acid desuccinylase DapE (123), which is involved in cell wall peptidoglycan production, the competence lipoprotein ComL (124), and the twin-arginine translocation (Tat) and general secretion (Sec) systems (125), which transport proteins across the cytoplasmic membrane (**Figure S9**). Furthermore, CP treatment resulted in downregulation of the oxidative stress sensing regulator *ospR* (126) and the long-chain fatty acid responsive regulator *psrA* (127) (**Figure S9**).

### CP elicits transcriptional responses indicative of redirected chorismate flux in *P. aeruginosa* cocultured with *S. aureus*

We also noticed a pattern amongst genes associated with the biosynthetic pathways utilizing chorismate, a key precursor for multiple secondary metabolites that contribute to the survival and virulence of *P. aeruginosa*. These secondary metabolites include the siderophore PCH (53, 128), the anthranilate-derived AQs (40), phenazines (129), the aromatic amino acid precursor prephenate (130), and 4-aminobenzoate (131) which is an intermediate for folate biosynthesis (132). DE analysis revealed that CP treatment, but not metal depletion, upregulated the expression of genes associated with the biosynthesis and utilization of 4-aminobenzoate and prephenate (**Figure 4** and **Table SF9**). These changes indicated increased cellular requirements for folate and amino acids which are needed to support metabolism. The expression of PCH biosynthetic machinery (133) was also upregulated as part of an Fe starvation response (*vide supra*) (68), indicating that some chorismate flux is directed towards PCH (**Figure 4**). CP treatment also significantly decreased the expression of genes encoding for phenazine biosynthetic machinery, which were among the most strongly downregulated genes identified (**Figures 4** and **S7**). These observations are in agreement with prior metabolite analyses of *P. aeruginosa* in monoculture and in coculture with *S. aureus* showing that Fe depletion significantly decreased phenazine levels and CP treatment resulted in near-complete suppression of phenazine production (59, 68). Moreover, the expression of *phnAB* anthranilate synthase (134–137) and AQ biosynthetic machinery (*pqs* operon) (42, 138) were downregulated in response to CP treatment; we observed that the effect of Fe limitation on these genes was attenuated as compared to CP treatment (**Figures 3** and **4**). Overall, our analysis uncovered that CP treatment elicits transcriptional responses indicative of redirected chorismate flux in *P. aeruginosa* in coculture with *S. aureus* and in monoculture (**Table SF9**). Our results also indicate that the transcriptional responses of *P. aeruginosa* to CP treatment are complex and involve both metal-dependent and metal-independent effects (68).

**Figure 4.**
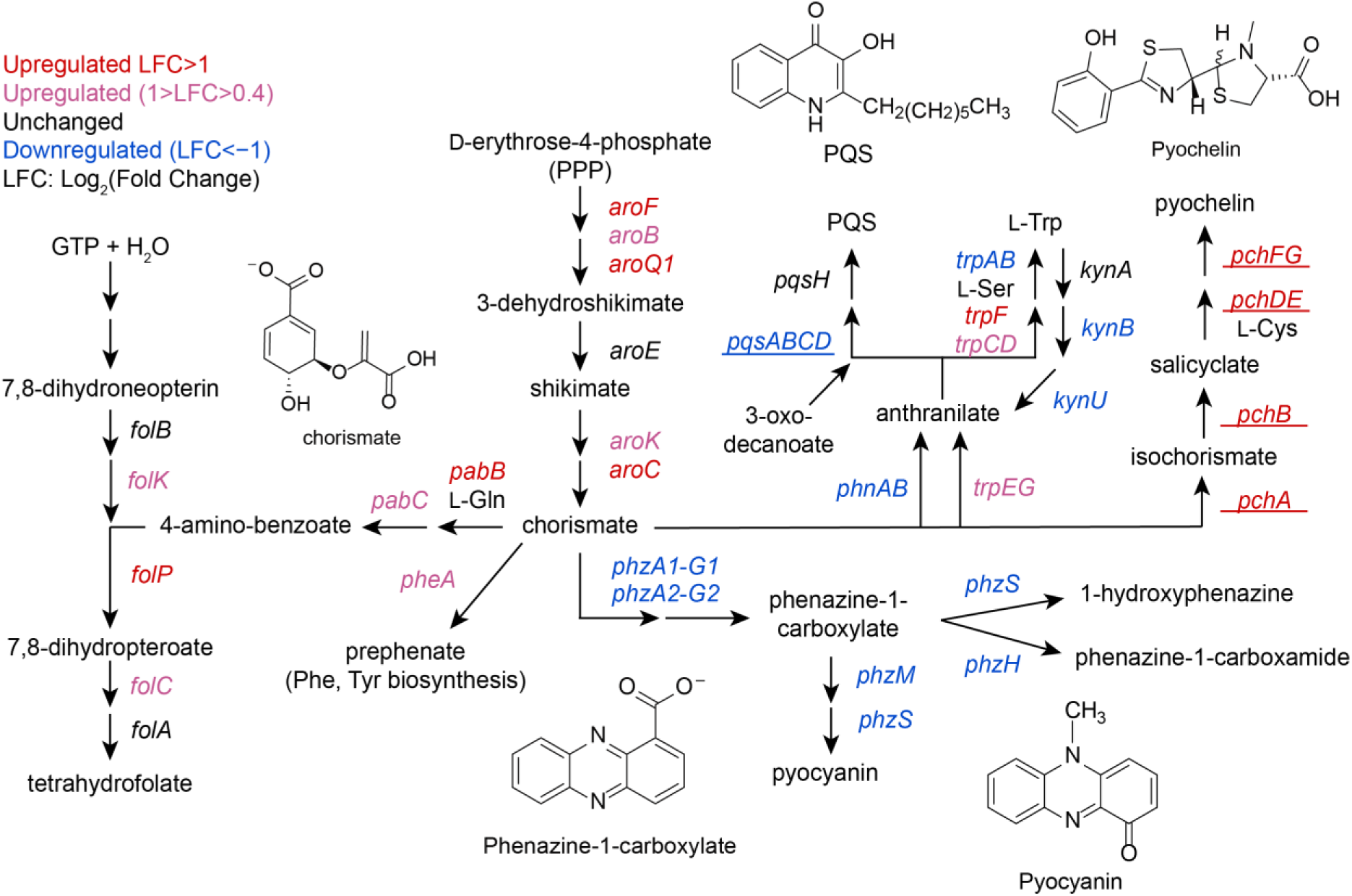
Gene expression analysis indicates that CP redirects chorismate flux in *P. aeruginosa* cocultured with *S. aureus*. *P. aeruginosa* genes DE in response to CP treatment are shown. Genes which were DE to a similar extent in CP-treated and Fe-depleted cocultures are underlined.

### CP perturbs autoinducer production and dampens quorum sensing by *P. aeruginosa* cocultured with *S. aureus*

DE analysis indicated that CP treatment resulted in the downregulation of *lasA*, *lasB*, and genes encoding for rhamnolipid production (*rhlAB*) (139) (**Figure 5A**), which are known to be controlled by QS. These trends were not observed in Fe-depleted or metal-depleted cocultures. To further interrogate these findings, we examined how CP treatment and Fe depletion affected the expression of these QS-controlled genes using real-time PCR. Consistent with trends revealed by RNA-seq, CP treatment decreased the expression of *lasA*, *lasB*, and *rhlA* for *P. aeruginosa* cocultured with *S. aureus* (**Figure 5B**). While Fe depletion slightly decreased the expression of *lasB*, this condition increased the expression of *lasA* and did not change the expression of *rhlA* (**Figure 5B**). These results indicate that CP attenuates the expression of QS-controlled genes for *P. aeruginosa* cocultured with *S. aureus* in a manner that is largely independent of Fe depletion.

**Figure 5.**
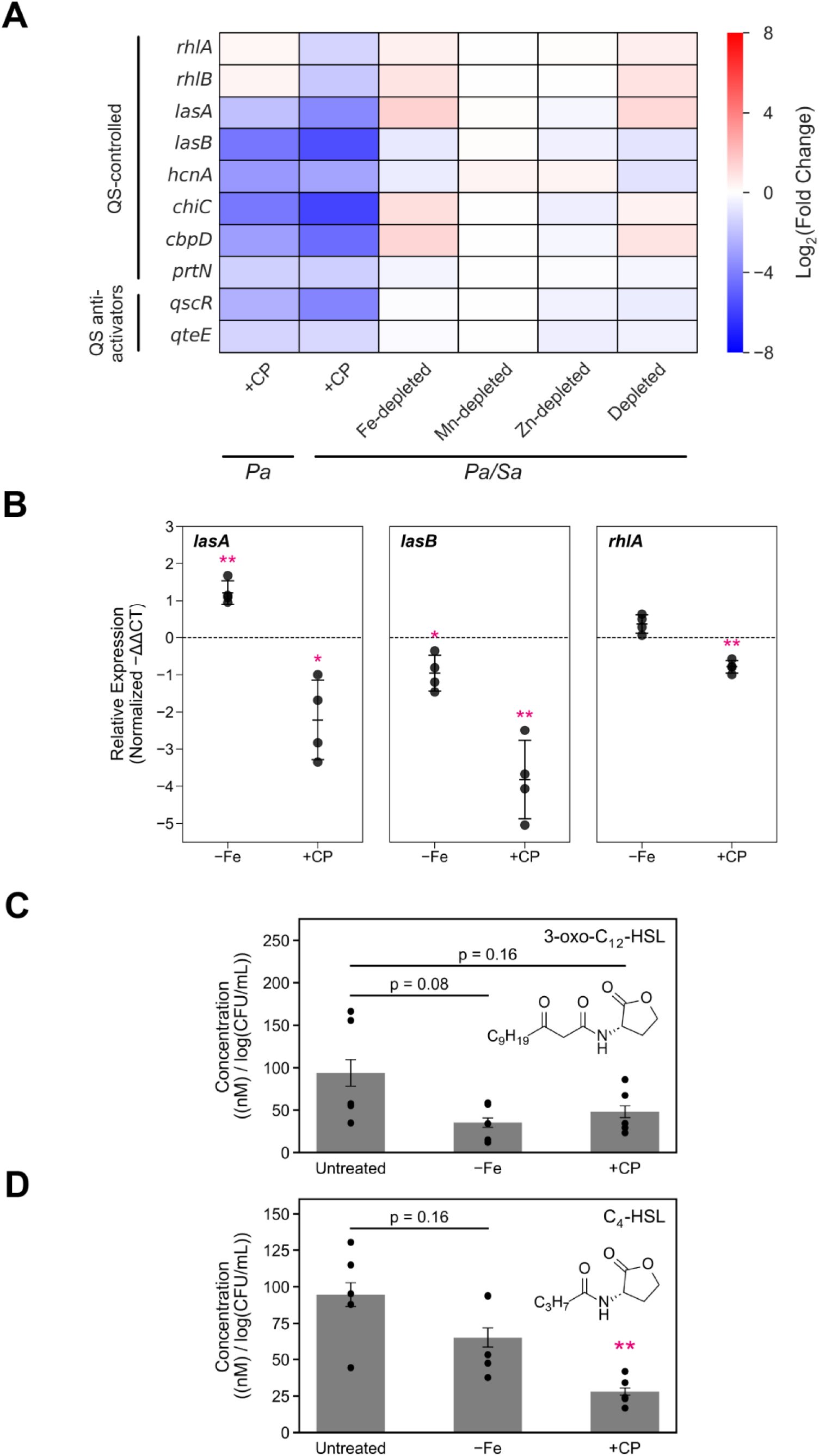
CP downregulates the expression of genes controlled by QS and decreases the production of autoinducers in *P. aeruginosa* cocultured with *S. aureus*. (**A**) DE heatmap of QS-controlled genes identified by RNA-seq. *Pa* indicates *P. aeruginosa* monoculture and *Pa/Sa* indicates the coculture. (**B**) DE of selected QS- controlled genes quantified by real-time PCR. (**C**, **D**) CP treatment and Fe depletion decrease the production of the QS autoinducers 3-oxo-C_12_-HSL (**C**) and C_4_-HSL (**D**) in *P. aeruginosa* cocultured with *S. aureus*. Cultures were grown in metal-replete CDM ± 20 μM CP or Fe-depleted CDM and incubated at 37 °C for 6 h. (**B**) Transcript levels were normalized to the *P. aeruginosa* housekeeping gene 16S and the fold change after normalization is presented (n=4, * p < 0.05, ** p < 0.01, error bars represent S.D.). (**C, D**) Aliquots were collected and processed for quantification by mass spectrometry. Metabolite levels were normalized to *P. aeruginosa* CFUs (n=5, error bars represent S.E.). For comparison with the untreated culture condition, ** p < 0.01.

Based on these findings, we suspected that CP may be acting on the LasIR and RhlIR QS systems, which govern regulation of LasA and LasB (22, 24, 140, 141). However, our efforts to understand the impact of CP on the expression of both QS systems were complicated by expression trends indicating overlap between the effects of CP treatment and Fe depletion. To further probe the effects of CP treatment on QS in *P. aeruginosa*, we utilized triple-quadrupole mass spectrometry to quantify levels of the autoinducers 3-oxo-C_12_-HSL and C_4_-HSL in coculture supernatants as a readout of QS molecules in *P. aeruginosa* (142, 143). Because CP treatment has negligible effects on the growth kinetics of *P. aeruginosa* cocultured with *S. aureus* (68), differences in levels of homoserine lactones (HSLs) are unlikely a result of differences in growth phase (144). Our mass spectrometric analysis revealed that CP treatment and Fe depletion slightly but insignificantly decreased levels of 3-oxo-C_12_-HSL (**Figure 5C**). By contrast, CP treatment markedly decreased levels of C_4_-HSL by approximately 70% at the 6 h timepoint (**Figure 5D**). Fe depletion caused no significant change in C_4_-HSL levels at the 6 h timepoint (**Figure 5D**). At the 11 h timepoint, CP treatment did not significantly alter levels of 3-oxo-C_12_-HSL (**Figure S10A**), indicating that production of 3-oxo-C_12_-HSL in CP-treated cocultures recovered to levels found in metal-replete cultures between 6 – 11 h. By contrast, levels of C_4_-HSL were decreased by both CP treatment and Fe depletion at 11 h (**Figure S10B**), indicating that the effect of CP on C_4_-HSL production occurs early in the culture time-course and persists through the 11 h timepoint. It is difficult to conclude from these data whether the effect of CP on C_4_-HSL levels can be attributed to Fe depletion.

To further understand how perturbed production of 3-oxo-C_12_-HSL and C_4_-HSL affected transcriptional responses of *P. aeruginosa*, we examined the expression of other *P. aeruginosa* genes which are known to be QS-controlled. Consistent with attenuated QS for *P. aeruginosa* in CP-treated cocultures, strong downregulation of genes encoding QS-controlled systems was also observed under these culture conditions. These genes encode phenazine biosynthetic machinery (*phz*), hydrogen cyanide production (*hcn*) (145), the chitin-binding protein CbpD (146), and PA-1 galactophilic lectin (*lecA*) (147, 148) (**Figures 5A and S9**). Furthermore, substantial overlap was found between genes that were DE in response to CP treatment and genes previously identified to be QS controlled by microarray analysis (24, 149). Out of 311 genes known to be positively regulated by QS (24), 245 (78.8%) were downregulated in CP-treated cocultures, of which 75 (24.1%) overlapped with the effects of Fe or Zn depletion and 170 (54.7%) were attributable only to the effects of CP treatment. We also observed that CP treatment resulted in downregulated expression of several genes encoding for recently identified QS anti-activator proteins, further suggesting perturbed QS (150). The presence of CP also led to decreased the expression of *qscR* (*PA14_39980*) (151), *qteE* (*PA14_30560*) (152) (**Figure 5A**), which is another indicator of perturbed QS. In addition, the QS-regulated *prtN* gene was downregulated in response to CP treatment (153) (**Figure 5A**). Collectively, these findings suggest that CP perturbs the production of *P. aeruginosa* AHL autoinducers primarily through effects on C_4_-HSL (**Figure 5D**), resulting in attenuated QS for *P. aeruginosa* in coculture. This noteworthy effect of CP on autoinducer production is likely to impact the expression of multiple virulence factors which directly impact coculture dynamics between *P. aeruginosa* and *S. aureus*.

### CP decreases the production of antimicrobials by *P. aeruginosa* cocultured with *S. aureus*

We recently showed that CP treatment and Fe depletion have opposing effects on *S. aureus* viability during coculture with *P. aeruginosa* (68). Previous and separate studies showed that CP treatment and Fe starvation also had opposing effects on AQ production (57, 66). Considering that these studies were done under different experimental conditions, we were motivated to evaluate the effects of CP treatment and Fe levels on AQ production by *P. aeruginosa* in parallel. *P. aeruginosa* produces multiple AQs such as 2-heptyl-4(1*H*)-quinolone (HHQ) (40–42, 58), 2-heptyl-3-hydroxy-4(1*H*)- quinolone (PQS) (40, 43), and the potent anti-staphylococcal metabolite 2-heptyl-4-hydroxyquinoline N-oxide (HQNO) (44). PQS and its precursor HHQ function as QS molecules that modulate activation of the virulence regulator MvfR (PqsR) and its regulon (40, 41, 154). PQS has been identified as a key QS signal molecule that controls many aspects of *P. aeruginosa* virulence (19, 155), including the expression of *lasB*. RNA-seq revealed that CP treatment decreased the expression of genes associated with AQ production (**Figures 3** and **4**). To further investigate these trends, we performed real-time PCR studies to examine the expression of AQ biosynthetic machinery in response to CP treatment and Fe depletion (42, 138). We observed that CP treatment slightly downregulated the expression of *pqsD* (**Figure S11A**) and decreased the expression of *pqsE* (**Figure S11B**), while Fe depletion resulted in significant downregulation of *pqsD* and *pqsE* (**Figure S11**), indicating that the effect of CP on *pqsD* and *pqsE* is attributable to Fe(II) sequestration by CP. Furthermore, we suspected that anthranilate production by *P. aeruginosa* may be reduced in the presence of CP due to downregulation of *phnAB* and *kynBU* (**Figure 4**), encoding the only two experimentally-verified sources of anthranilate for AQ production (42, 135, 156–158).

Given transcriptional responses indicating decreased AQ production in response to CP treatment, we used mass spectrometry to ascertain whether CP treatment decreased the production of AQs by *P. aeruginosa* cocultured with *S. aureus*. The alkyl chains in AQs can vary in length and saturation with the C_7_ congeners being the first studied (20, 159, 160). Because *P. aeruginosa* also synthesizes C_9_ AQ congeners (41) which possess anti-staphylococcal activity similar to the C_7_ congeners (58), we included the C_9_ AQ congeners, 2-nonyl-4(*1H*)-quinolone (NHQ), 2-nonyl-4-hydroxyquinoline N-oxide (NQNO), and 2-nonyl-3-hydroxy-4(*1H*)-quinolone (C_9_-PQS) in our analysis. Using triple-quadrupole mass spectrometry, we quantified AQ levels from the same coculture supernatants used for the analysis of HSLs. Strikingly, CP treatment considerably decreased levels of HHQ, HQNO and PQS at the 6 h (**Figures S12A – S12C**) and 11 h (**Figures 6A – 6C**) timepoints. While Fe depletion also decreased levels of HHQ at both time points, levels of HQNO were increased by Fe depletion at both timepoints (**Figures S12A**, **S12B**, **6A** and **6B**), although we note that in some cases the difference was not statistically significant. PQS levels in Fe-depleted cocultures were slightly increased at 6 h (**Figure S12C**) and decreased at 11 h (**Figure 6C**), suggesting that production of PQS in cocultures grown in Fe-depleted CDM may be negatively autoregulated at this later time point during growth. CP treatment and Fe depletion decreased the overall pool of C_7_-AQs at both timepoints (**Figures 6D** and **S13**), which was primarily driven by decreased levels of HHQ.

**Figure 6.**
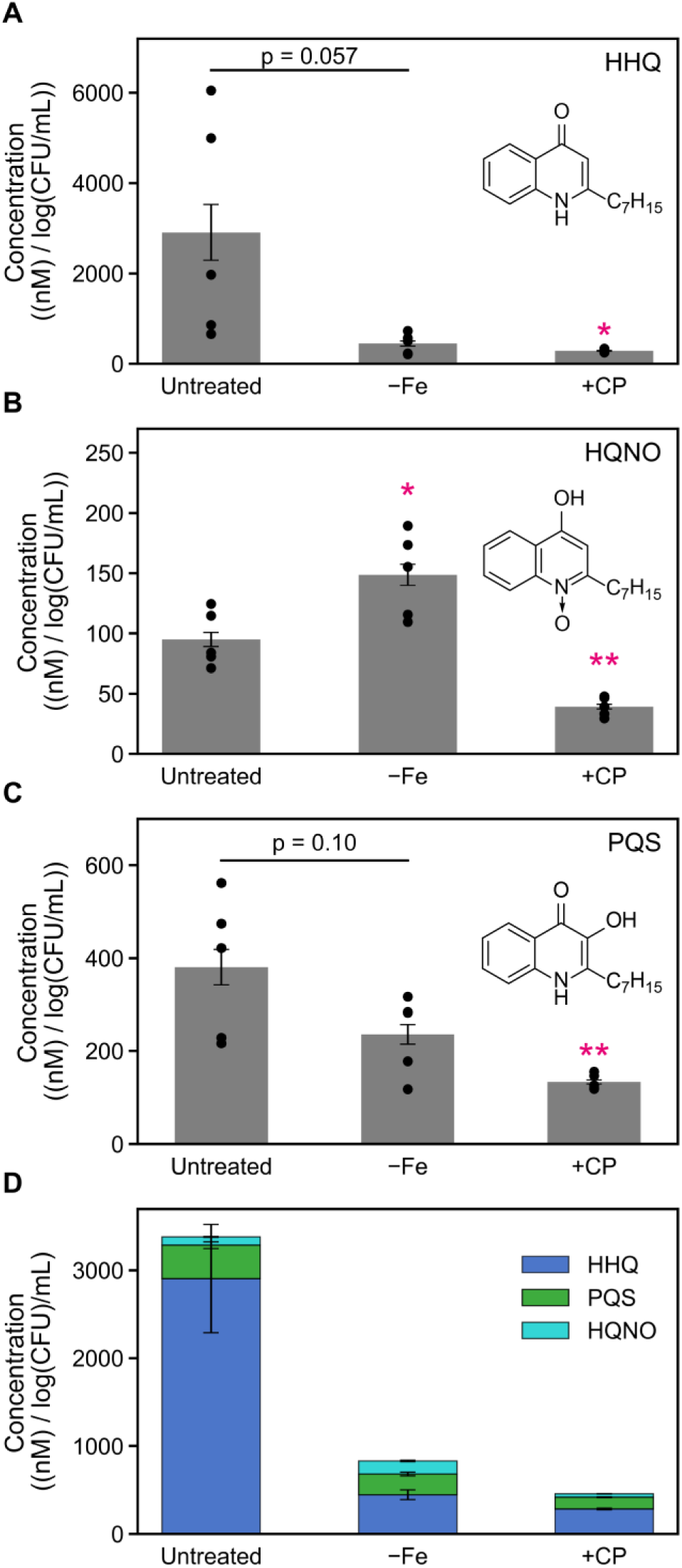
The presence of CP decreases the production of C_7_ AQs in *P. aeruginosa*/*S. aureus* cocultures. CP treatment decreases the levels of HHQ (**A**), HQNO (**B**), PQS (**C**) and the overall C_7_ AQ pool (**D**). Aliquots were collected from cocultures of *P. aeruginosa* and *S. aureus* grown in Fe-depleted CDM or metal-replete CDM ± 20 μM CP at 37 °C, 11 h, and processed for quantification by mass spectrometry. Metabolite levels were normalized to *P. aeruginosa* CFUs (n=5, error bars represent S.E.). For comparison with the untreated culture condition, * p < 0.05, ** p < 0.01.

Consistent with the C_7_ congeners, CP treatment resulted in a marked decrease in NHQ levels at both the 6 h (**Figure S14A**) and 11 h (**Figure 7A**) timepoints, a decrease that was not observed for Fe-depleted cocultures. By contrast, NQNO levels were significantly increased in Fe-depleted cocultures and unchanged in CP-treated cocultures at both timepoints (**Figures S14B** and **7B**). Levels of C_9_-PQS were increased in CP-treated cocultures and Fe-depleted cocultures at the 6 h timepoint **(Figure S14C**), but were not significantly altered at 11 h (**Figure 7C**), in agreement with trends observed for C_7_-PQS (**Figures S12C** and **6C**). Changes in the overall pool of C_9_ AQs were primarily driven by changes in the levels of C_9_-PQS (**Figures S15** and **7D**). Together, our findings show that the effect of CP on the C_9_ AQs occurs primarily on NHQ, which exhibits significant anti-staphylococcal activity in low Fe conditions (58) such as those expected in the presence of CP.

**Figure 7.**
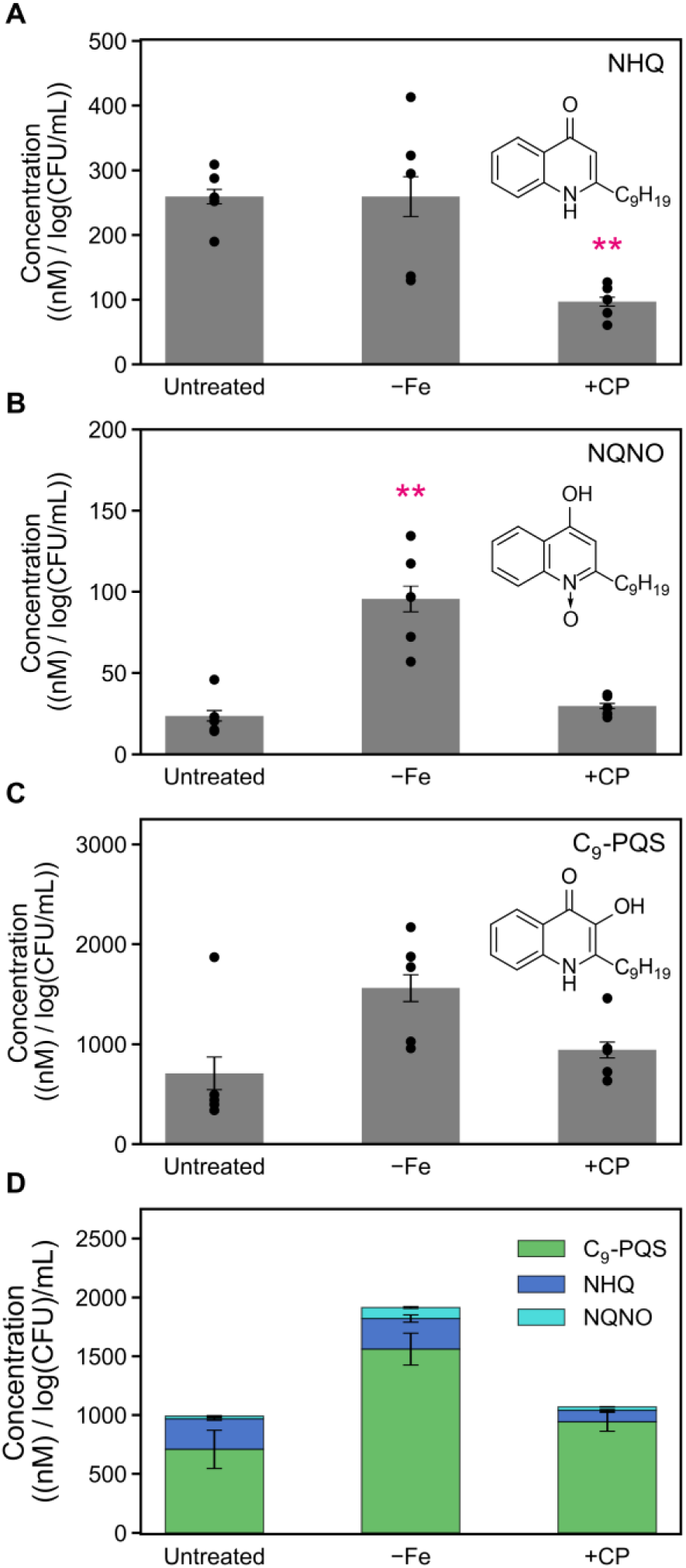
The presence of CP decreases levels of the C_9_ AQ NHQ in *P. aeruginosa*/*S. aureus* co-cultures. CP treatment decreased the levels of NHQ (**A**) but did not significantly affect levels of NQNO (**B**), C_9_- PQS (**C**) or the overall C_9_ AQ pool (**D**). Aliquots were collected from cocultures of *P. aeruginosa* and *S. aureus* grown in Fe-depleted CDM or metal-replete CDM ± 20 μM CP at 37 °C, 11 h, and processed for quantification by mass spectrometry. Metabolite levels were normalized to *P. aeruginosa* CFUs (n=5, error bars represent S.E.). For comparison with the untreated culture condition, ** p < 0.01.

Overall, our analysis of AQ production by *P. aeruginosa* in coculture is consistent with the transcriptional responses indicating redirected chorismate flux, with decreased chorismate flux towards phenazine and AQ production (**Figure 4**). Furthermore, our results are in agreement with prior observations of the opposing effects of CP treatment and Fe depletion on *S. aureus* survival in coculture with *P. aeruginosa* (57, 58, 66, 68). Collectively, our findings indicate that one aspect of CP-mediated *S. aureus* survival in coculture with *P. aeruginosa* (66, 68) involves decreased production of the C_7_ AQs HHQ, HQNO and PQS, and the C_9_ AQ NHQ. The observation that CP treatment decreased PQS production was also consistent with CP-induced downregulation of *psrA* (161) (**Figure S9**), a gene known to positively regulate PQS production. By contrast, *P. aeruginosa* produced increased levels of the anti- staphylococcal metabolites HQNO and NQNO in Fe-depleted cocultures, which contributes towards heightened anti-staphylococcal activity by *P. aeruginosa* in Fe-limited environments.

### *S. aureus* cocultured with *P. aeruginosa* mounts Fe starvation responses in the presence of CP but not in Fe-depleted conditions

The above analysis highlights the distinct impacts of CP treatment and Fe limitation on *P. aeruginosa* AQ production, with likely consequences on *S. aureus* viability in coculture. To gain improved understanding of how *S. aureus* responds to CP treatment and metal limitation, analysis of the *S. aureus* transcriptome during coculture was performed. In response to CP, *S. aureus* upregulated multiple systems associated with metal acquisition and siderophore utilization (**Figures 8A** and **8C**), indicating that CP elicits multi-metal starvation responses from *S. aureus* in coculture. Our findings are consistent with prior studies examining the transcriptional responses of *S. aureus* to CP treatment, which include real-time PCR studies of *S. aureus* cocultured with *P. aeruginosa* (68) and a recent RNA-seq study of *S. aureus* monocultures (62). However, Venn analysis of the top 300 DE genes across all conditions revealed only partial overlap between the transcriptional responses of *S. aureus* in coculture to CP treatment and metal depletion (**Figures 8A** and **8B**), prompting further analysis of these distinct effects (**Figures S16 – S20**).

**Figure 8.**
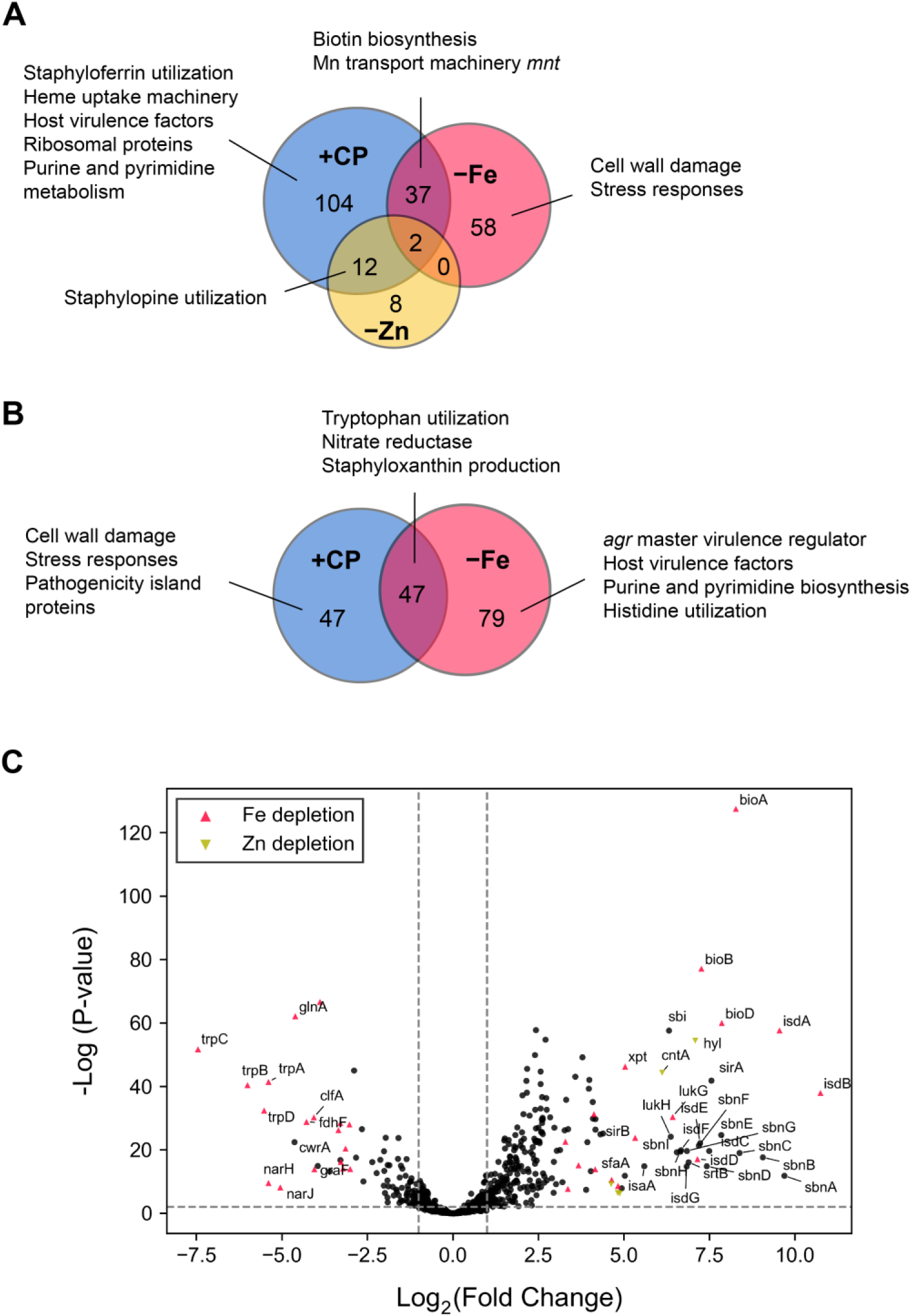
DE profiles of CP treatment and Fe depletion overlap in *S. aureus* cocultured with *P. aeruginosa*. Venn diagrams of the top 600 DE *S. aureus* genes across all conditions tested reveals partial overlap for upregulated (**A**) genes, and considerable overlap of downregulated (**B**) genes in CP treated cocultures and cocultures grown in Fe-depleted CDM or metal-depleted CDM. (**C**) Volcano plot of DE changes in response to CP treatment. Genes with similar DE patterns in response to Fe depletion or Zn depletion are denoted as colored shapes. A threshold cutoff Log_2_(Fold Change) of 1 was employed.

In agreement with prior qPCR studies, CP treatment elicited robust Fe starvation responses from *S. aureus* in monoculture (61, 62) and in coculture with *P. aeruginosa*. These responses include the upregulation of genes encoding for heme uptake machinery (*isd*) (56) and staphyloferrin biosynthesis and transport (*sir* and *sbn*) (162, 163). Unexpectedly, the upregulation of these hallmark Fe starvation responses was not observed for *S. aureus* in Fe-depleted cocultures and was attenuated for *S. aureus* in metal-depleted cocultures (**Figure 9**). These surprising results suggest that the ability of *S. aureus* to mount Fe starvation responses is inhibited by the antistaphylococcal activity of *P. aeruginosa*, which is exacerbated by Fe depletion and mitigated by CP treatment (57, 58, 66, 68).

**Figure 9.**
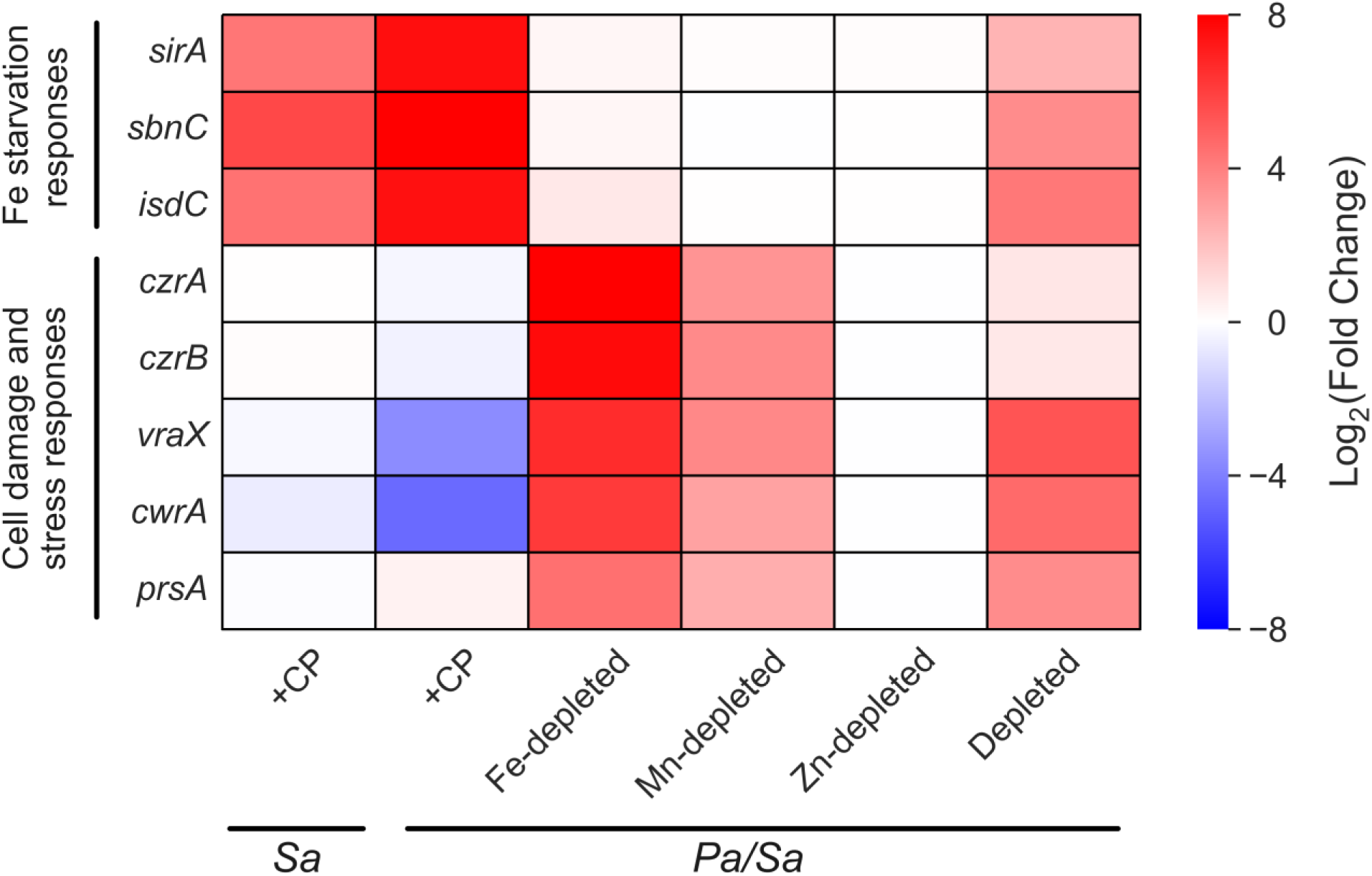
The presence of CP elicits Fe starvation responses and decreases cell damage and stress responses by *S. aureus* cocultured with *P. aeruginosa*. DE heatmap of *S. aureus* genes associated with Fe starvation responses and cell damage and stress responses. *Sa* indicates *S. aureus* monoculture and *Pa/Sa* indicates the coculture.

In agreement with this notion, a cluster of genes was found to be upregulated in Fe-depleted cocultures but not in CP-treated cocultures, which contained the Zn-responsive transcriptional repressor and efflux transporter *czrAB* (164, 165), the cell wall inhibition responsive protein *cwrA* (166), the peptidylprolyl isomerase *prsA* (167, 168), and superoxide stress responses (**Figures 9** and **S21**), reflecting increased levels of cell wall stress and damage in *S. aureus* in Fe-depleted cocultures. Transcriptional responses of *S. aureus* in metal-depleted cocultures indicated decreased severity of cell wall damage and stress as compared to Fe-depleted cocultures (**Figures 9** and **S21**) despite depletion of either Fe or all three metals having similar impacts on *S. aureus* viability (68). CP treatment also resulted in transcriptional responses indicating increased translational activity, evident from the upregulation of genes associated with translational machinery and metabolism, and downregulation of the genes encoding for the transcriptional repressor LexA (169, 170) and the ribosome-associated inhibitor protein RaiA (171) (**Figure S22**). By contrast, genes associated with translational activity were downregulated for *S. aureus* in Fe-depleted cocultures; this downregulation was found to be attenuated for *S. aureus* in metal-depleted cocultures (**Figure S22**).

These data point to a model in which CP promotes the survival of *S. aureus* in coculture with *P. aeruginosa* by reducing the anti-staphylococcal activity of *P. aeruginosa*, as evident from transcriptional responses indicating perturbed QS and decreased AQ production in *P. aeruginosa*, and decreased cell wall damage and increased metabolism in *S. aureus*. As a result, *S. aureus* cocultured with *P. aeruginosa* in the presence of CP is able to mount Fe starvation responses. We speculate that the ability of *S*. *aureus* to mount Fe starvation responses, although attenuated, in metal-depleted cocultures but not in Fe-depleted cocultures may stem from effects of multi-metal depletion (**Supplementary Discussion**).

### CP increases the expression of genes associated with host virulence in *S. aureus* cocultured with *P. aeruginosa*

CP upregulated the expression of multiple *S. aureus* systems asssociated with host virulence in coculture, in agreement with a prior RNA-seq study examining the effect of CP on *S. aureus* Newman in monoculture (62). We observed that CP uniquely upregulated the expression of genes encoding for the immunoglobin-binding protein Sbi (172), alpha-hemolysin (*hyl*) (173), lytic transglycoysylase IsaA (174), the secretory antigen SsaA (175), and the phenol-soluble modulins (*psm*) (176) in coculture (**Figure 10** and **Table SF19**). These findings were consistent with CP-induced upregulation of the *saeRS* two-component system (177–179), which regulates many *S. aureus* host virulence factors (**Figure 10**).

**Figure 10.**
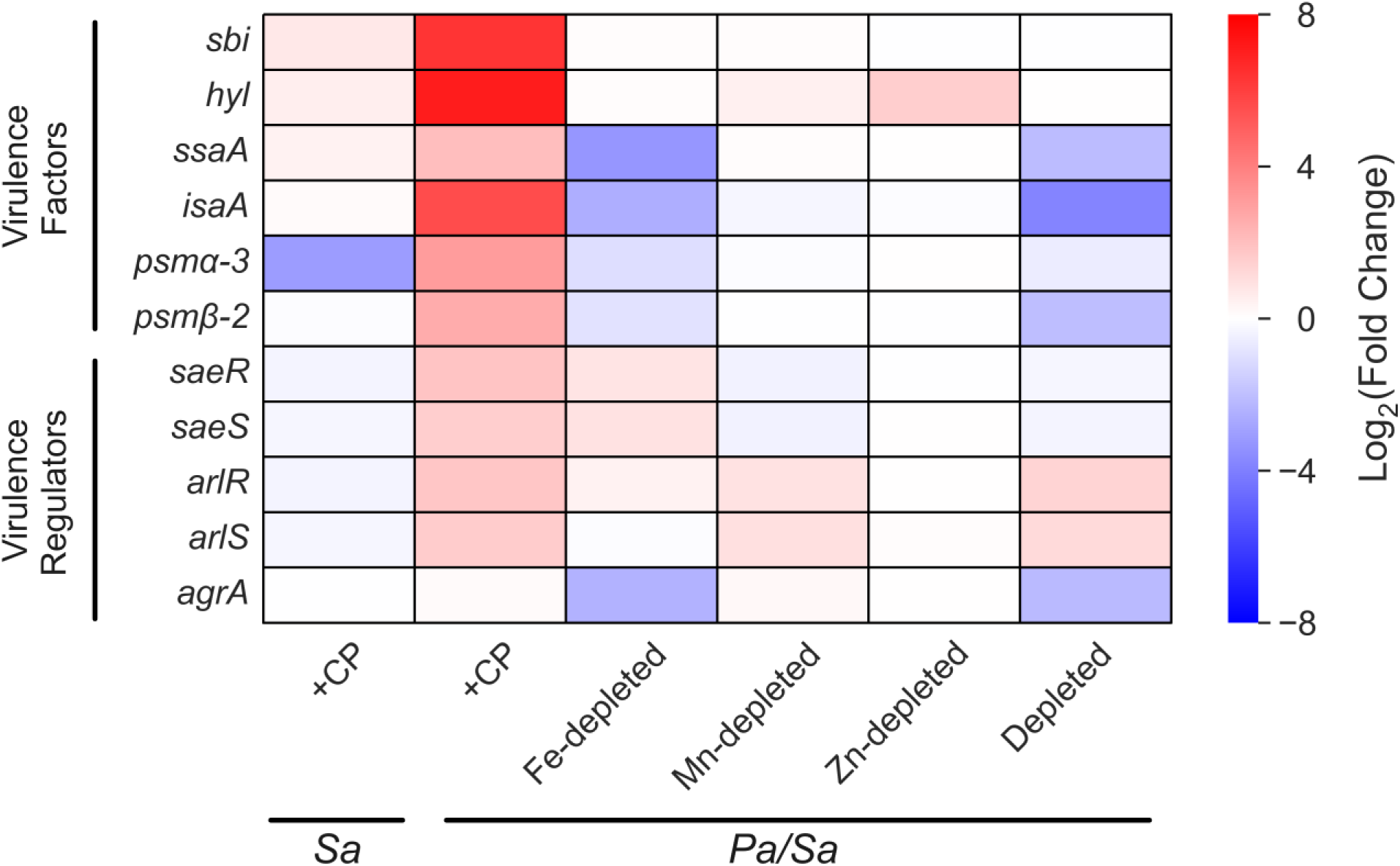
CP upregulates the expression of virulence factors and regulators in *S. aureus* cocultured with *P. aeruginosa*. DE heatmap of *S. aureus* genes associated with host virulence factors and virulence regulation. *Sa* indicates *S. aureus* monoculture and *Pa/Sa* indicates the coculture.

Expression of the *agr* master virulence regulator (180) remained unchanged in response to CP treatment and was decreased in response to Fe depletion (**Table**s **SF15B** and **SF20**). In addition, expression of the *S. aureus* QS effector RNAIII was decreased in response to CP treatment, Fe depletion, and Mn depletion (**Figure S23**). Together, these findings indicate that CP-mediated upregulation of *S. aureus* host virulence genes is unlikely due to *agr*-mediated regulation or metal sequestration by CP. The upregulation of *psmα- 3* was consistent with increased expression of the sRNA Teg41 (*srna_1080_RsaX05*) (**Figure S23** and **Table SF21**), which was previously shown to be important for virulence in *S. aureus* and required for the expression of the alpha phenol-soluble modulins (181, 182). Intriguingly, CP increased the expression of the sRNA *rsaA* for *S. aureus* in coculture (**Figure S23**), which is known to suppress the translation of the pleiotropic virulence regulator MgrA (183, 184). We note that the effect of CP on *S. aureus* monocultures was attenuated (**Figure 10**), which most likely stems from variations in the growth phase of *S. aureus* and culture conditions between this work and previous investigations (62, 68).

We also observed that CP treatment resulted in upregulated expression of the *dltABCD* operon (62, 185) for *S. aureus* in coculture, which is responsible for the modication of cell wall teichoic acids in response to environmental stresses (**Figure S24**). The presence of CP also led to upregulation of the *kdpDE* two-component system (186), which is involved in virulence regulation by sensing extracellular K(I) levels. In addition, CP downregulated the expression of genes encoding for Clp protease, which is involved in protein homeostasis and the expression of various virulence factors in *S. aureus* (62, 187). Lastly, we observed that the presence of CP slightly perturbed the expression of two global regulators, *sigB* (188, 189) and *sarA* (190–192), which are involved in *S. aureus* virulence and adaptation (**Figure S24**). We did not observe comparable upregulation of *S. aureus* host virulence genes in monocultures treated with CP. Overall, the increased expression of host virulence genes in the presence of CP is consistent with the protective effect of CP on *S. aureus* cocultured with *P. aeruginosa*, and highlights the profound impact of this protection on the *S. aureus* transcriptome.

### Working Model and Outlook

The insights discussed here provide a working model for how CP and Fe availability impact coculture dynamics between *P. aeruginosa* and *S. aureus* (**Figure 11A**). In this model, *P. aeruginosa* functions as an attacker that produces an arsenal of anti-staphylococcal factors, while *S. aureus* functions as a defender. Under metal-replete and metal-depleted conditions, *P. aeruginosa* attacks *S. aureus* via AQNOs and other secreted factors, and the inability of *S. aureus* to defend itself leads to decreased viability. CP effectively disarms *P. aeruginosa* by perturbing autoinducer production, redirecting chorismate flux, and decreasing the production of AQs. By decreasing the anti-staphylococcal activity of *P. aeruginosa*, the presence of CP increases the viability of *S. aureus* cocultured with *P. aeruginosa* and thus *S. aureus* mounts Fe starvation and host virulence responses that are otherwise not feasible in the presence of *P. aeruginosa*. Our findings also highlight complexity in the transcriptional responses of both bacterial pathogens to CP, some of which overlap with responses to metal depletion and others of which appear to result from metal-independent effects of the protein.

**Figure 11.**
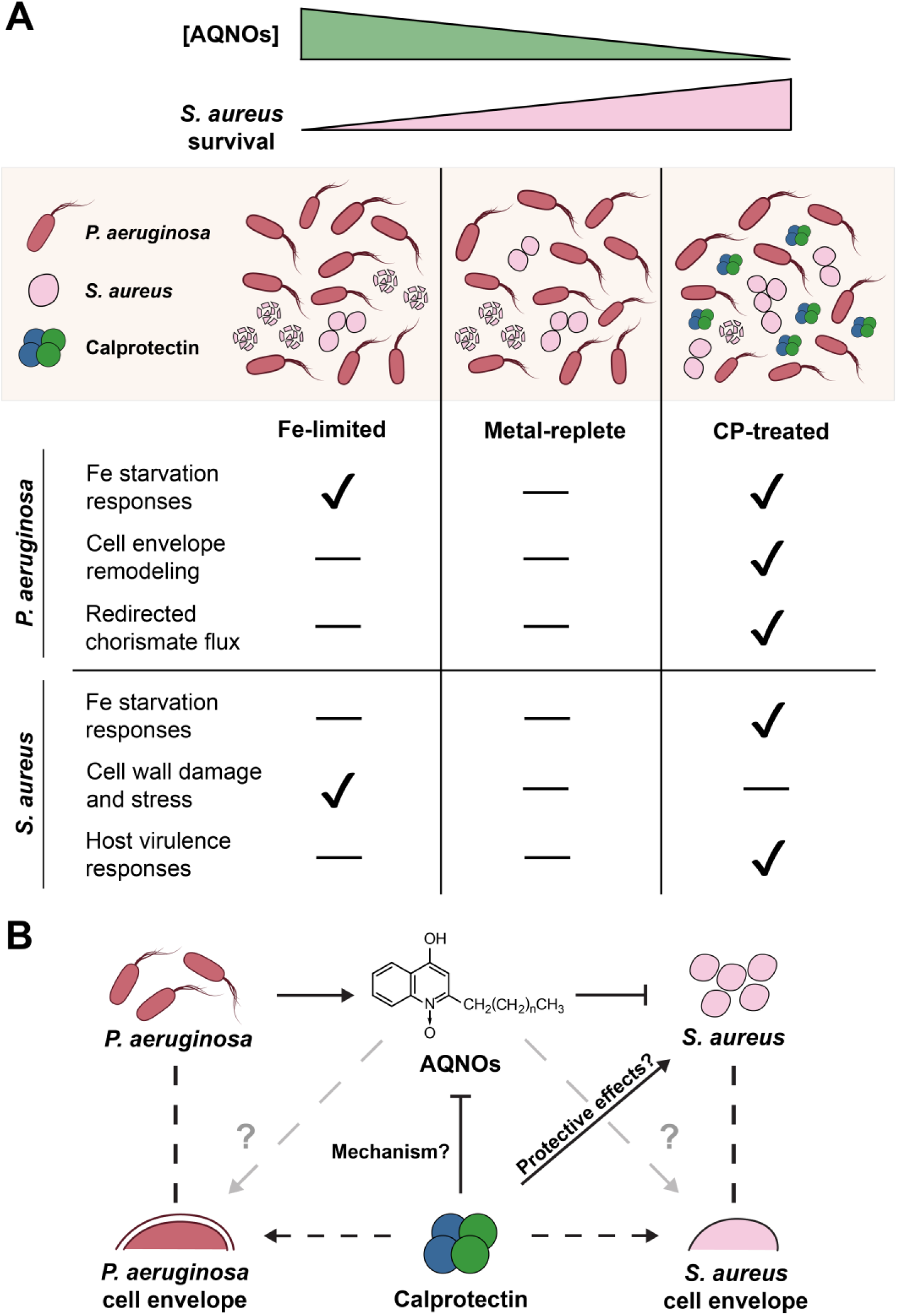
Current working model of the impact of CP on coculture dynamics between *P. aeruginosa* and *S. aureus*. (**A**) CP treatment and Fe depletion have distinct and opposing impacts on coculture dynamics. (**B**) Scheme of interactions between CP, AQNOs, and cocultures of *P. aeruginosa* and *S. aureus*.

This study and the resulting model present several outstanding questions that warrant future investigation (**Figure 11B)**. Firstly, this work motivates investigating the mechanism by which AQ levels are modulated by CP treatment, i.e., does CP directly initiate a *P. aeruginosa* signaling cascade that results in decreased AQ production, or does *P. aeruginosa* reduce AQ production due to a distinct physiological response to CP? Secondly, the collective results from prior studies (60, 62, 68) and this work point to CP eliciting cell envelope changes in *P. aeruginosa* and *S. aureus*. Whether these membrane changes are linked to the observed decrease in AQ production remains to be investigated. Thirdly, future work should examine whether CP-mediated protection of *S. aureus* results from CP boosting *S. aureus* defenses against *P. aeruginosa* – perhaps through cell envelope modifications – or if coculture dynamics are primarily driven by changes in AQ production. Fourthly, given that CP-mediated protection of *S. aureus* occurs independently of metal sequestration, future studies should determine whether the CP protein scaffold, absent its metal-binding sites, can recapitulate the transcriptional responses elicited by CP from cocultures of *P. aeruginosa* and *S*. *aureus* (68), or if any other structural features or functions of CP are required. Lastly, we expect this model to pertain to the extracellular host environment where CP is released from the neutrophil and encounters these two bacterial pathogens, and the impact of these interactions on the host warrants exploration. These insights will likely be key to understanding the multifaceted activity of this remarkable protein and its consequences on interspecies interactions at the host–pathogen interface, such as in polymicrobial infections.

## Materials and Methods

For complete materials and methods, please refer to the accompanying Supplementary Information.

### General experimental methods

General methods, including general microbiology methods and protein production and handling, were performed as reported in (68).

### RNA extraction and workup for RNA-seq

RNA extraction and workup was carried out as previously reported (68). Following RNA precipitation and resuspension in nuclease-free water, samples were submitted to the MIT BioMicro Center for further preparation and sequencing. Sample integrity was validated using fluorescence-based electrophoresis (AATI Fragment Analyzer), and ribosomal depletion was performed using the NEB Next® rRNA Depletion Kit (New England Biolabs). Subsequently, library preparation (adapter ligation, size selection, barcoding, enrichment) was performed using the NEBNext Ultra II Directional RNA Library Prep Kit (New England Biolabs). The quality of the resulting libraries was validated using real-time PCR, following which the libraries were pooled and sequenced on a single lane of an Illumina NextSeq500 instrument using 75 nt chemistry.

### Bioinformatics workflow and analyses

Raw reads were aligned with the hisat2 aligner (193) using NCBI RefSeq reference genomes for *P. aeruginosa* PA14 (GCF_000014625.1) (194) and *S. aureus JE2* (GCF_002085525.1) (Walter Reed Army Institute of Research). The combined genome derived from the PA14 + JE2 genomes was used to align reads from coculture samples, and mapping fidelity was validated by verifying no cross-mapping occurred. The aligned reads were quantified using Feature Aggregate Depth Utility (195). To check for sequencing depth and the presence of technical artifacts, rarefaction analysis and principal component analysis were performed. DE analysis of the quantified reads was performed using DESeq2 (version 1.40.2) (196). For both monocultures and cocultures, untreated cultures (in metal-replete CDM) served as the untreated control. Log2(fold changes) (LFCs) were calculated using the apeglm method for effect size shrinkage (197). Functional enrichment analyses were performed with clusterProfiler (198, 199) using the variance-stabilized LFCs for both species against the best available annotations for each species. To access these annotations, cross-species gene mapping was carried out with CD-HIT-EST-2D (200) to obtain unique gene mappings. For *P. aeruginosa*, overrepresentation analysis was performed using curated PAO1 GO annotation available on the *Pseudomonas* genome database (pseudomonas.com) (201). For *S. aureus*, gene set enrichment analysis was performed using the NCBI Refseq annotations available for *S. aureus* USA300 FPR3757 (CP000255.1) (202).

### Metabolite quantification by triple quadrupole mass spectrometry

HPLC-grade solvents were used for sample preparation and mass spectrometry. A 350 μL aliquot of culture suspension was centrifuged at 13,000 rpm, 5 min, 4 °C to pellet cells and debris. A 300 μL aliquot of the supernatant was transferred to a new polypropylene tube and 3 μL of 100 μM C_6_-HSL-d_3_ internal standard in methanol was added. The resulting mixture was extracted twice with an equal volume of acidified ethyl acetate containing 0.02% v/v acetic acid, each time by vigorous vortexing at 3000 rpm, ambient temperature for 1 min. The upper organic layers were transferred to a clean glass vial and the solvent was removed by rotatory evaporation in a 35 °C water bath. The resulting solid was resuspended in 900 μL of ice-cold methanol and transferred to a new ice-cold tube. The resuspended samples were centrifuged at 13,000 rpm, 10 min, 4 °C to pellet any particulates, and 200 μL of the supernatant was transferred into a HPLC vial fitted with a polypropylene vial insert for analysis. Complete instrumentation data and HPLC conditions are provided in the Supplementary Information.

### Preparation of analyte standards

The homoserine lactones and alkylquinolone standards were obtained from commercial vendors and used as received. To determine the dynamic range of detection and obtain standard curves, analyte standards (2 nM – 50 μM) were prepared by serial dilution of fresh 1 – 10 mM stocks of each analyte in methanol, with the exception of C_9_-PQS which was instead dissolved into a 1:1 mixture of water:acetonitrile, each containing 0.1% trifluoroacetic acid. The analyte standards were centrifuged at 13,000 rpm, 10 min, 4 °C to pellet any particulates and loaded into vials as described above.

## Acknowledgements

This work was supported by the NIH (R01 GM126376 to EMN and AGO). W.H.L. received support from the A*STAR National Science Scholarship (BS-PhD). The triple quadrupole MS instrument is housed and maintained in the MIT DCIF (HHMI). Real-time PCR and RNA-seq instrumentation are housed and maintained in the MIT BioMicro Center (NIH P30-ES002109). The authors acknowledge Dr. Mohanraja Kumar (MIT DCIF) for helpful discussions regarding mass spectrometry, Dr. Stuart Levine (MIT BioMicro Center) for helpful recommendations for RNA-seq, Dr. Charlie Whittaker (MIT Swanson Biotechnology Center) for assistance with CD-HIT-EST and initial processing of raw RNA-seq reads, and Dr. Vincent Butty (MIT BioMicro Center), Prof. Alex Shalek (MIT), and Prof. Julie Hotopp (Univ. of Maryland) for helpful RNA-seq discussions. The authors thank Jonathon Gans for technical assistance with real-time PCR primers for *crcZ*.

## References

1. Ciofu O, Tolker-Nielsen T. 2019. Tolerance and Resistance of *Pseudomonas aeruginosa* Biofilms to Antimicrobial Agents—How *Pseudomonas aeruginosa* Can Escape Antibiotics. Front Microbiol 10.

2. Howden BP, Giulieri SG, Wong Fok Lung T, Baines SL, Sharkey LK, Lee JYH, Hachani A, Monk IR, Stinear TP. 2023. *Staphylococcus aureus* host interactions and adaptation. Nat Rev Microbiol 21:380–395.

3. De Oliveira DMP, Forde BM, Kidd TJ, Harris PNA, Schembri MA, Beatson SA, Paterson DL, Walker MJ. 2020. Antimicrobial Resistance in ESKAPE Pathogens. Clin Microbiol Rev 33:e00181–19.

4. Camus L, Briaud P, Vandenesch F, Doléans-Jordheim A, Moreau K. 2022. Mixed Populations and Co-Infection: *Pseudomonas aeruginosa* and *Staphylococcus aureus*. Adv Exp Med Biol 1386:397– 424.

5. DeLeon S, Clinton A, Fowler H, Everett J, Horswill AR, Rumbaugh KP. 2014. Synergistic interactions of *Pseudomonas aeruginosa* and *Staphylococcus aureus* in an *in vitro* wound model. Infect Immun 82:4718–4728.

6. Sibley CD, Duan K, Fischer C, Parkins MD, Storey DG, Rabin HR, Surette MG. 2008. Discerning the Complexity of Community Interactions Using a *Drosophila* Model of Polymicrobial Infections. PLOS Pathogens 4:e1000184.

7. Korgaonkar A, Trivedi U, Rumbaugh KP, Whiteley M. 2013. Community surveillance enhances *Pseudomonas aeruginosa* virulence during polymicrobial infection. Proc Natl Acad Sci U S A 110:1059–1064.

8. Alves PM, Al-Badi E, Withycombe C, Jones PM, Purdy KJ, Maddocks SE. 2018. Interaction between *Staphylococcus aureus* and *Pseudomonas aeruginosa* is beneficial for colonisation and pathogenicity in a mixed biofilm. Pathog Dis 76:fty003.

9. Dalton T, Dowd SE, Wolcott RD, Sun Y, Watters C, Griswold JA, Rumbaugh KP. 2011. An In Vivo Polymicrobial Biofilm Wound Infection Model to Study Interspecies Interactions. PLoS ONE 6:e27317.

10. Michelsen CF, Christensen A-MJ, Bojer MS, Høiby N, Ingmer H, Jelsbak L. 2014. *Staphylococcus aureus* Alters Growth Activity, Autolysis, and Antibiotic Tolerance in a Human Host-Adapted *Pseudomonas aeruginosa* Lineage. J Bacteriol 196:3903–3911.

11. Orazi G, O’Toole GA. 2017. *Pseudomonas aeruginosa* Alters *Staphylococcus aureus* Sensitivity to Vancomycin in a Biofilm Model of Cystic Fibrosis Infection. mBio 8:e00873–17.

12. Tognon M, Köhler T, Gdaniec BG, Hao Y, Lam JS, Beaume M, Luscher A, Buckling A, van Delden C. 2017. Co-evolution with *Staphylococcus aureus* leads to lipopolysaccharide alterations in *Pseudomonas aeruginosa*. ISME J 11:2233–2243.

13. Smith AC, Rice A, Sutton B, Gabrilska R, Wessel AK, Whiteley M, Rumbaugh KP. 2017. Albumin inhibits *Pseudomonas aeruginosa* quorum sensing and alters polymicrobial interactions. Infect Immun 85:e00116–17.

14. Nguyen AT, Oglesby-Sherrouse AG. 2016. Interactions between *Pseudomonas aeruginosa* and *Staphylococcus aureus* during co-cultivations and polymicrobial infections. Appl Microbiol Biotechnol 100:6141–6148.

15. Filkins LM, Graber JA, Olson DG, Dolben EL, Lynd LR, Bhuju S, O’Toole GA. 2015. Coculture of *Staphylococcus aureus* with *Pseudomonas aeruginosa* drives *S. aureus* towards fermentative metabolism and reduced viability in a cystic fibrosis model. J Bacteriol 197:2252–2264.

16. Hotterbeekx A, Kumar-Singh S, Goossens H, Malhotra-Kumar S. 2017. *In vivo* and *in vitro* interactions between *Pseudomonas aeruginosa* and *Staphylococcus* spp. Front Cell Infect Microbiol 7:106.

17. Pierson LS, Pierson EA. 2010. Metabolism and function of phenazines in bacteria: impacts on the behavior of bacteria in the environment and biotechnological processes. Appl Microbiol Biotechnol 86:1659–1670.

18. Pearson JP, Pesci EC, Iglewski BH. 1997. Roles of *Pseudomonas aeruginosa las* and *rhl* quorum-sensing systems in control of elastase and rhamnolipid biosynthesis genes. J Bacteriol 179:5756– 5767.

19. Pesci EC, Pearson JP, Seed PC, Iglewski BH. 1997. Regulation of *las* and *rhl* quorum sensing in *Pseudomonas aeruginosa*. J Bacteriol 179:3127–3132.

20. Pesci EC, Milbank JBJ, Pearson JP, McKnight S, Kende AS, Greenberg EP, Iglewski BH. 1999. Quinolone signaling in the cell-to-cell communication system of *Pseudomonas aeruginosa*. Proc Natl Acad Sci U S A 96:11229–11234.

21. Brint JM, Ohman DE. 1995. Synthesis of multiple exoproducts in *Pseudomonas aeruginosa* is under the control of RhlR-RhlI, another set of regulators in strain PAO1 with homology to the autoinducer-responsive LuxR-LuxI family. J Bacteriol 177:7155–7163.

22. Anderson RM, Zimprich CA, Rust L. 1999. A Second Operator Is Involved in *Pseudomonas aeruginosa* Elastase (*lasB*) Activation. J Bacteriol 181:6264–6270.

23. Winson MK, Camara M, Latifi A, Foglino M, Chhabra SR, Daykin M, Bally M, Chapon V, Salmond GP, Bycroft BW. 1995. Multiple N-acyl-L-homoserine lactone signal molecules regulate production of virulence determinants and secondary metabolites in *Pseudomonas aeruginosa*. Proc Natl Acad Sci U S A 92:9427–9431.

24. Schuster M, Lostroh CP, Ogi T, Greenberg EP. 2003. Identification, Timing, and Signal Specificity of *Pseudomonas aeruginosa* Quorum-Controlled Genes: a Transcriptome Analysis. J Bacteriol 185:2066–2079.

25. Latifi A, Foglino M, Tanaka K, Williams P, Lazdunski A. 1996. A hierarchical quorum-sensing cascade in *Pseudomonas aeruginosa* links the transcriptional activators LasR and RhIR (VsmR) to expression of the stationary-phase sigma factor RpoS. Mol Microbiol 21:1137–1146.

26. Pearson JP, Van Delden C, Iglewski BH. 1999. Active efflux and diffusion are involved in transport of *Pseudomonas aeruginosa* cell-to-cell signals. J Bacteriol 181:1203–1210.

27. Whiteley M, Lee KM, Greenberg EP. 1999. Identification of genes controlled by quorum sensing in *Pseudomonas aeruginosa*. Proc Natl Acad Sci U S A 96:13904.

28. Pearson JP, Gray KM, Passador L, Tucker KD, Eberhard A, Iglewski BH, Greenberg EP. 1994. Structure of the autoinducer required for expression of *Pseudomonas aeruginosa* virulence genes. Proc Natl Acad Sci U S A 91:197–201.

29. Pearson JP, Passador L, Iglewski BH, Greenberg EP. 1995. A second N-acylhomoserine lactone signal produced by *Pseudomonas aeruginosa*. Proc Natl Acad Sci U S A 92:1490–1494.

30. Gambello MJ, Iglewski BH. 1991. Cloning and characterization of the *Pseudomonas aeruginosa lasR* gene, a transcriptional activator of elastase expression. J Bacteriol 173:3000–3009.

31. Passador L, Cook JM, Gambello MJ, Rust L, Iglewski BH. 1993. Expression of *Pseudomonas aeruginosa* virulence genes requires cell-to-cell communication. Science 260:1127–1130.

32. Seed PC, Passador L, Iglewski BH. 1995. Activation of the *Pseudomonas aeruginosa lasI* gene by LasR and the *Pseudomonas* autoinducer PAI: an autoinduction regulatory hierarchy. J Bacteriol 177:654–659.

33. Toder DS, Gambello MJ, Iglewski BH. 1991. *Pseudomonas aeruginosa* LasA: a second elastase under the transcriptional control of *lasR*. Mol Microbiol 5:2003–2010.

34. Kessler E, Safrin M, Olson JC, Ohman DE. 1993. Secreted LasA of *Pseudomonas aeruginosa* is a staphylolytic protease. J Biol Chem 268:7503–7508.

35. Kessler E, Safrin M, Gustin JK, Ohman DE. 1998. Elastase and the LasA Protease of *Pseudomonas aeruginosa* Are Secreted with Their Propeptides. J Biol Chem 273:30225–30231.

36. Bever RA, Iglewski BH. 1988. Molecular characterization and nucleotide sequence of the *Pseudomonas aeruginosa* elastase structural gene. J Bacteriol 170:4309–4314.

37. Braun P, de Groot A, Bitter W, Tommassen J. 1998. Secretion of elastinolytic enzymes and their propeptides by *Pseudomonas aeruginosa*. J Bacteriol 180:3467–3469.

38. Kessler E, Safrin M. 1988. Synthesis, processing, and transport of *Pseudomonas aeruginosa* elastase. J Bacteriol 170:5241–5247.

39. Dietrich LEP, Price-Whelan A, Petersen A, Whiteley M, Newman DK. 2006. The phenazine pyocyanin is a terminal signalling factor in the quorum sensing network of *Pseudomonas aeruginosa*. Mol Microbiol 61:1308–1321.

40. Diggle SP, Matthijs S, Wright VJ, Fletcher MP, Chhabra SR, Lamont IL, Kong X, Hider RC, Cornelis P, Cámara M, Williams P. 2007. The *Pseudomonas aeruginosa* 4-Quinolone Signal Molecules HHQ and PQS Play Multifunctional Roles in Quorum Sensing and Iron Entrapment. Chem Biol 14:87–96.

41. Déziel E, Lépine F, Milot S, He J, Mindrinos MN, Tompkins RG, Rahme LG. 2004. Analysis of *Pseudomonas aeruginosa* 4-hydroxy-2-alkylquinolines (HAQs) reveals a role for 4-hydroxy-2- heptylquinoline in cell-to-cell communication. Proc Natl Acad Sci U S A 101:1339–1344.

42. Gallagher LA, McKnight SL, Kuznetsova MS, Pesci EC, Manoil C. 2002. Functions required for extracellular quinolone signaling by *Pseudomonas aeruginosa*. J Bacteriol 184:6472–6480.

43. Pesci EC, Milbank JBJ, Pearson JP, McKnight S, Kende AS, Greenberg EP, Iglewski BH. 1999. Quinolone signaling in the cell-to-cell communication system of Pseudomonas aeruginosa. Proc Natl Acad Sci U S A 96:11229–11234.

44. Machan ZA, Taylor GW, Pitt TL, Cole PJ, Wilson R. 1992. 2-Heptyl-4-hydroxyquinoline N-oxide, an antistaphylococcal agent produced by *Pseudomonas aeruginosa*. J Antimicrob Chemother 30:615–623.

45. Hoffman LR, Déziel E, D’Argenio DA, Lépine F, Emerson J, McNamara S, Gibson RL, Ramsey BW, Miller SI. 2006. Selection for *Staphylococcus aureus* small-colony variants due to growth in the presence of *Pseudomonas aeruginosa*. Proc Natl Acad Sci U S A 103:19890–19895.

46. Voggu L, Schlag S, Biswas R, Rosenstein R, Rausch C, Götz F. 2006. Microevolution of cytochrome bd oxidase in Staphylococci and its implication in resistance to respiratory toxins released by *Pseudomonas*. J Bacteriol 188:8079–8086.

47. Murdoch CC, Skaar EP. 2022. Nutritional immunity: the battle for nutrient metals at the host– pathogen interface. 11. Nat Rev Microbiol 20:657–670.

48. Zygiel EM, Nolan EM. 2018. Transition metal sequestration by the host-defense protein calprotectin. Annu Rev Biochem 87:621–643.

49. Cartron ML, Maddocks S, Gillingham P, Craven CJ, Andrews SC. 2006. Feo – transport of ferrous iron into bacteria. Biometals 19:143–157.

50. Lau CKY, Krewulak KD, Vogel HJ. 2016. Bacterial ferrous iron transport: the Feo system. FEMS Microbiology Reviews 40:273–298.

51. Wendenbaum S, Demange P, Dell A, Meyer JM, Abdallah MA. 1983. The structure of pyoverdine Pa, the siderophore of *Pseudomonas aeruginosa*. Tetrahedron Letters 24:4877–4880.

52. Visca P, Imperi F, Lamont IL. 2007. Pyoverdine siderophores: from biogenesis to biosignificance. Trends Microbiol 15:22–30.

53. Cox CD, Rinehart KL, Moore ML, Cook JC. 1981. Pyochelin: novel structure of an iron-chelating growth promoter for *Pseudomonas aeruginosa*. Proc Natl Acad Sci U S A 78:4256–4260.

54. Ghssein G, Brutesco C, Ouerdane L, Fojcik C, Izaute A, Wang S, Hajjar C, Lobinski R, Lemaire D, Richaud P, Voulhoux R, Espaillat A, Cava F, Pignol D, Borezée-Durant E, Arnoux P. 2016. Biosynthesis of a broad-spectrum nicotianamine-like metallophore in *Staphylococcus aureus*. Science 352:1105–1109.

55. Smith AD, Wilks A. 2015. Differential contributions of the outer membrane receptors PhuR and HasR to heme acquisition in *Pseudomonas aeruginosa*. J Biol Chem 290:7756–7766.

56. Skaar EP, Schneewind O. 2004. Iron-regulated surface determinants (Isd) of *Staphylococcus aureus*: stealing iron from heme. Microbes and Infection 6:390–397.

57. Nguyen AT, Jones JW, Ruge MA, Kane MA, Oglesby-Sherrouse AG. 2015. Iron depletion enhances production of antimicrobials by *Pseudomonas aeruginosa*. J Bacteriol 197:2265–2275.

58. Nguyen AT, Jones JW, Cámara M, Williams P, Kane MA, Oglesby-Sherrouse AG. 2016. Cystic fibrosis isolates of *Pseudomonas aeruginosa* retain iron-regulated antimicrobial activity against *Staphylococcus aureus* through the action of multiple alkylquinolones. Front Microbiol 7:1171.

59. Zygiel EM, Nelson CE, Brewer LK, Oglesby-Sherrouse AG, Nolan EM. 2019. The human innate immune protein calprotectin induces iron starvation responses in *Pseudomonas aeruginosa*. J Biol Chem 294:3549–3562.

60. Nelson CE, Huang W, Zygiel EM, Nolan EM, Kane MA, Oglesby AG. 2021. The human innate immune protein calprotectin elicits a multimetal starvation response in *Pseudomonas aeruginosa*. Microbiol Spectr 9:e00519–21.

61. Zygiel EM, Obisesan AO, Nelson CE, Oglesby AG, Nolan EM. 2021. Heme protects *Pseudomonas aeruginosa* and *Staphylococcus aureus* from calprotectin-induced iron starvation. J Biol Chem 296:100160.

62. Reyes Ruiz VM, Freiberg JA, Weiss A, Green ER, Jobson M-E, Felton E, Shaw LN, Chazin WJ, Skaar EP. 2024. Coordinated adaptation of *Staphylococcus aureus* to calprotectin-dependent metal sequestration. mBio 15:e01389–24.

63. Kehl-Fie TE, Chitayat S, Hood MI, Damo S, Restrepo N, Garcia C, Munro KA, Chazin WJ, Skaar EP. 2011. Nutrient metal sequestration by calprotectin inhibits bacterial superoxide defense enhancing neutrophil killing of *Staphylococcus aureus*. Cell Host Microbe 10:158–164.

64. Damo SM, Kehl-Fie TE, Sugitani N, Holt ME, Rathi S, Murphy WJ, Zhang Y, Betz C, Hench L, Fritz G, Skaar EP, Chazin WJ. 2013. Molecular basis for manganese sequestration by calprotectin and roles in the innate immune response to invading bacterial pathogens. Proc Natl Acad Sci U S A 110:3841–3846.

65. Besold AN, Gilston BA, Radin JN, Ramsoomair C, Culbertson EM, Li CX, Cormack BP, Chazin WJ, Kehl-Fie TE, Culotta VC. 2018. Role of calprotectin in withholding zinc and copper from *Candida albicans*. Infect Immun 86:e00779–17.

66. Wakeman CA, Moore JL, Noto MJ, Zhang Y, Singleton MD, Prentice BM, Gilston BA, Doster RS, Gaddy JA, Chazin WJ, Caprioli RM, Skaar EP. 2016. The innate immune protein calprotectin promotes *Pseudomonas aeruginosa* and *Staphylococcus aureus* interaction. Nat Commun 7:11951.

67. Nakashige TG, Zhang B, Krebs C, Nolan EM. 2015. Human calprotectin is an iron-sequestering host-defense protein. Nat Chem Biol 11:765–771.

68. Lee WH, Zygiel EM, Lee, CH, Oglesby AG, Nolan EM. 2025. Calprotectin-mediated survival of *Staphylococcus aureus* in coculture with *Pseudomonas aeruginosa* occurs without nutrient metal sequestration. mBio 10.1128/mBio.03846-24R1.

69. Baishya J, Everett JA, Chazin WJ, Rumbaugh KP, Wakeman CA. 2022. The innate immune protein calprotectin interacts with and encases biofilm communities of *Pseudomonas aeruginosa* and *Staphylococcus aureus*. Front Cell Infect Microbiol 12:898796.

70. Gray RD, Imrie M, Boyd AC, Porteous D, Innes JA, Greening AP. 2010. Sputum and serum calprotectin are useful biomarkers during CF exacerbation. J Cyst Fibros 9:193–198.

71. Reid DW, Lam QT, Schneider H, Walters EH. 2004. Airway iron and iron-regulatory cytokines in cystic fibrosis. Eur Respir J 24:286–291.

72. Johne B, Fagerhol MK, Lyberg T, Prydz H, Brandtzaeg P, Naess-Andresen CF, Dale I. 1997. Functional and clinical aspects of the myelomonocyte protein calprotectin. Mol Pathol 50:113–123.

73. Herrera OR, Christensen ML, Helms RA. 2016. Calprotectin: clinical applications in pediatrics. J Pediatr Pharmacol Ther 21:308–321.

74. Dixon P. 2003. VEGAN, a package of R functions for community ecology. J Veg Sci 14:927–930.

75. Hassett DJ, Schweizer HP, Ohman DE. 1995. *Pseudomonas aeruginosa sodA* and *sodB* mutants defective in manganese- and iron-cofactored superoxide dismutase activity demonstrate the importance of the iron-cofactored form in aerobic metabolism. J Bacteriol 177:6330–6337.

76. Létoffé S, Redeker V, Wandersman C. 1998. Isolation and characterization of an extracellular haem-binding protein from *Pseudomonas aeruginosa* that shares function and sequence similarities with the *Serratia marcescens* HasA haemophore. Mol Microbiol 28:1223–1234.

77. Iglewski BH, Kabat D. 1975. NAD-dependent inhibition of protein synthesis by *Pseudomonas aeruginosa* toxin. Proc Natl Acad Sci U S A 72:2284–2288.

78. Hedstrom RC, Funk CR, Kaper JB, Pavlovskis OR, Galloway DR. 1986. Cloning of a gene involved in regulation of exotoxin A expression in *Pseudomonas aeruginosa*. Infect Immun 51:37– 42.

79. Gambello MJ, Kaye S, Iglewski BH. 1993. LasR of *Pseudomonas aeruginosa* is a transcriptional activator of the alkaline protease gene (*apr*) and an enhancer of exotoxin A expression. Infect Immun 61:1180–1184.

80. Nelson CE, Huang W, Brewer LK, Nguyen AT, Kane MA, Wilks A, Oglesby-Sherrouse AG. 2019. Proteomic Analysis of the *Pseudomonas aeruginosa* Iron Starvation Response Reveals PrrF Small Regulatory RNA-Dependent Iron Regulation of Twitching Motility, Amino Acid Metabolism, and Zinc Homeostasis Proteins. J Bacteriol 201:e00754–18.

81. Lamarche MG, Déziel E. 2011. MexEF-OprN Efflux Pump Exports the *Pseudomonas* Quinolone Signal (PQS) Precursor HHQ (4-hydroxy-2-heptylquinoline). PLoS ONE 6:e24310.

82. Köhler T, van Delden C, Curty LK, Hamzehpour MM, Pechere JC. 2001. Overexpression of the MexEF-OprN multidrug efflux system affects cell-to-cell signaling in *Pseudomonas aeruginosa*. J Bacteriol 183:5213–5222.

83. Fetar H, Gilmour C, Klinoski R, Daigle DM, Dean CR, Poole K. 2010. *mexEF-oprN* Multidrug Efflux Operon of *Pseudomonas aeruginosa*: Regulation by the MexT Activator in Response to Nitrosative Stress and Chloramphenicol. Antimicrob Agents Chemother 55:508.

84. Hassett DJ, Howell ML, Ochsner UA, Vasil ML, Johnson Z, Dean GE. 1997. An operon containing *fumC* and *sodA* encoding fumarase C and manganese superoxide dismutase is controlled by the ferric uptake regulator in *Pseudomonas aeruginosa*: *fur* mutants produce elevated alginate levels. J Bacteriol 179:1452.

85. Wilderman PJ, Sowa NA, FitzGerald DJ, FitzGerald PC, Gottesman S, Ochsner UA, Vasil ML. 2004. Identification of tandem duplicate regulatory small RNAs in *Pseudomonas aeruginosa* involved in iron homeostasis. Proc Natl Acad Sci U S A 101:9792–9797.

86. Oglesby-Sherrouse AG, Vasil ML. 2010. Characterization of a Heme-Regulated Non-Coding RNA Encoded by the *prrF* Locus of *Pseudomonas aeruginosa*. PLoS ONE 5:e9930.

87. Reinhart AA, Powell DA, Nguyen AT, O’Neill M, Djapgne L, Wilks A, Ernst RK, Oglesby-Sherrouse AG. 2015. The *prrF*-encoded small regulatory RNAs are required for iron homeostasis and virulence of *Pseudomonas aeruginosa*. Infect Immun 83:863–875.

88. Heo Y-J, Chung I-Y, Cho W-J, Lee B-Y, Kim J-H, Choi K-H, Lee J-W, Hassett DJ, Cho Y-H. 2010. The Major Catalase Gene (*katA*) of *Pseudomonas aeruginosa* PA14 Is under both Positive and Negative Control of the Global Transactivator OxyR in Response to Hydrogen Peroxide. J Bacteriol 192:381–390.

89. Su S, Panmanee W, Wilson JJ, Mahtani HK, Li Q, VanderWielen BD, Makris TM, Rogers M, McDaniel C, Lipscomb JD, Irvin RT, Schurr MJ, Lancaster JR, Kovall RA, Hassett DJ. 2014. Catalase (KatA) Plays a Role in Protection against Anaerobic Nitric Oxide in *Pseudomonas aeruginosa*. PLoS ONE 9:e91813.

90. Hassett DJ, Woodruff WA, Wozniak DJ, Vasil ML, Cohen MS, Ohman DE. 1993. Cloning and characterization of the *Pseudomonas aeruginosa sodA* and *sodB* genes encoding manganese- and iron-cofactored superoxide dismutase: demonstration of increased manganese superoxide dismutase activity in alginate-producing bacteria. J Bacteriol 175:7658–7665.

91. Mandrand-Berthelot MA, Couchoux-Luthaud G, Santini CL, Giordano G. 1988. Mutants of *Escherichia coli* specifically deficient in respiratory formate dehydrogenase activity. J Gen Microbiol 134:3129–3139.

92. Schlindwein C, Giordano G, Santini CL, Mandrand MA. 1990. Identification and expression of the *Escherichia coli fdhD* and *fdhE* genes, which are involved in the formation of respiratory formate dehydrogenase. J Bacteriol 172:6112–6121.

93. Borrero-de Acuña JM, Rohde M, Wissing J, Jänsch L, Schobert M, Molinari G, Timmis KN, Jahn M, Jahn D. 2016. Protein Network of the *Pseudomonas aeruginosa* Denitrification Apparatus. J Bacteriol 198:1401–1413.

94. Comolli JC, Donohue TJ. 2004. Differences in two *Pseudomonas aeruginosa cbb* cytochrome oxidases. Mol Microbiol 51:1193–1203.

95. Kawakami T, Kuroki M, Ishii M, Igarashi Y, Arai H. 2010. Differential expression of multiple terminal oxidases for aerobic respiration in *Pseudomonas aeruginosa*. Environ Microbiol 12:1399– 1412.

96. Arai H, Kawakami T, Osamura T, Hirai T, Sakai Y, Ishii M. 2014. Enzymatic Characterization and In Vivo Function of Five Terminal Oxidases in *Pseudomonas aeruginosa*. J Bacteriol 196:4206– 4215.

97. Weidner U, Geier S, Ptock A, Friedrich T, Leif H, Weiss H. 1993. The gene locus of the proton-translocating NADH: ubiquinone oxidoreductase in *Escherichia coli*. Organization of the 14 genes and relationship between the derived proteins and subunits of mitochondrial complex I. J Mol Biol 233:109–122.

98. Yao H, Jepkorir G, Lovell S, Nama PV, Weeratunga S, Battaile KP, Rivera M. 2011. Two distinct ferritin-like molecules in *Pseudomonas aeruginosa*: the product of the *bfrA* gene is a bacterial ferritin (FtnA) and not a bacterioferritin (Bfr). Biochemistry 50:5236–5248.

99. Lhospice S, Gomez NO, Ouerdane L, Brutesco C, Ghssein G, Hajjar C, Liratni A, Wang S, Richaud P, Bleves S, Ball G, Borezée-Durant E, Lobinski R, Pignol D, Arnoux P, Voulhoux R. 2017. *Pseudomonas aeruginosa* zinc uptake in chelating environment is primarily mediated by the metallophore pseudopaline. Sci Rep 7:17132.

100. Mastropasqua MC, D’Orazio M, Cerasi M, Pacello F, Gismondi A, Canini A, Canuti L, Consalvo A, Ciavardelli D, Chirullo B, Pasquali P, Battistoni A. 2017. Growth of *Pseudomonas aeruginosa* in zinc poor environments is promoted by a nicotianamine-related metallophore. Mol Microbiol 106:543–561.

101. Patzer SI, Hantke K. 2000. The zinc-responsive regulator Zur and its control of the *znu* gene cluster encoding the ZnuABC zinc uptake system in *Escherichia coli*. J Biol Chem 275:24321–24332.

102. Pederick VG, Eijkelkamp BA, Begg SL, Ween MP, McAllister LJ, Paton JC, McDevitt CA. 2015. ZnuA and zinc homeostasis in *Pseudomonas aeruginosa*. Sci Rep 5:13139.

103. Tsui HC, Zhao G, Feng G, Leung HC, Winkler ME. 1994. The mutL repair gene of *Escherichia coli* K-12 forms a superoperon with a gene encoding a new cell-wall amidase. Mol Microbiol 11:189– 202.

104. Scheurwater EM, Pfeffer JM, Clarke AJ. 2007. Production and purification of the bacterial autolysin N-acetylmuramoyl-L-alanine amidase B from *Pseudomonas aeruginosa*. Protein Expr Purif 56:128–137.

105. Blaby-Haas CE, Furman R, Rodionov DA, Artsimovitch I, de Crécy-Lagard V. 2011. Role of a Zn-independent DksA in Zn homeostasis and stringent response. Mol Microbiol 79:700–715.

106. Furman R, Biswas T, Danhart EM, Foster MP, Tsodikov OV, Artsimovitch I. 2013. DksA2, a zinc-independent structural analog of the transcription factor DksA. FEBS Letters 587:614–619.

107. De Bentzmann S, Giraud C, Bernard CS, Calderon V, Ewald F, Plésiat P, Nguyen C, Grunwald D, Attree I, Jeannot K, Fauvarque M-O, Bordi C. 2012. Unique Biofilm Signature, Drug Susceptibility and Decreased Virulence in *Drosophila* through the *Pseudomonas aeruginosa* Two-Component System PprAB. PLoS Pathog 8:e1003052.

108. Kammler M, Schön C, Hantke K. 1993. Characterization of the ferrous iron uptake system of *Escherichia coli*. J Bacteriol 175:6212–6219.

109. Nau CD, Konisky J. 1989. Evolutionary relationship between the TonB-dependent outer membrane transport proteins: nucleotide and amino acid sequences of the *Escherichia coli* colicin I receptor gene. J Bacteriol 171:1041–1047.

110. Rabsch W, Methner U, Voigt W, Tschäpe H, Reissbrodt R, Williams PH. 2003. Role of Receptor Proteins for Enterobactin and 2,3-Dihydroxybenzoylserine in Virulence of *Salmonella enterica*. Infect Immun 71:6953.

111. Johnson L, Mulcahy H, Kanevets U, Shi Y, Lewenza S. 2012. Surface-localized spermidine protects the *Pseudomonas aeruginosa* outer membrane from antibiotic treatment and oxidative stress. J Bacteriol 194:813–826.

112. Smith EE, Buckley DG, Wu Z, Saenphimmachak C, Hoffman LR, D’Argenio DA, Miller SI, Ramsey BW, Speert DP, Moskowitz SM, Burns JL, Kaul R, Olson MV. 2006. Genetic adaptation by *Pseudomonas aeruginosa* to the airways of cystic fibrosis patients. Proc Natl Acad Sci U S A 103:8487–8492.

113. Gunn JS, Lim KB, Krueger J, Kim K, Guo L, Hackett M, Miller SI. 1998. PmrA-PmrB-regulated genes necessary for 4-aminoarabinose lipid A modification and polymyxin resistance. Mol Microbiol 27:1171–1182.

114. Gunn JS, Ryan SS, Van Velkinburgh JC, Ernst RK, Miller SI. 2000. Genetic and functional analysis of a PmrA-PmrB-regulated locus necessary for lipopolysaccharide modification, antimicrobial peptide resistance, and oral virulence of *Salmonella enterica* serovar Typhimurium. Infect Immun 68:6139–6146.

115. de Bentzmann S, Aurouze M, Ball G, Filloux A. 2006. FppA, a novel *Pseudomonas aeruginosa* prepilin peptidase involved in assembly of type IVb pili. J Bacteriol 188:4851–4860.

116. Bernard CS, Bordi C, Termine E, Filloux A, de Bentzmann S. 2009. Organization and PprB- Dependent Control of the *Pseudomonas aeruginosa tad* Locus, Involved in Flp Pilus Biology. J Bacteriol 191:1961–1973.

117. Giraud C, Bernard CS, Calderon V, Yang L, Filloux A, Molin S, Fichant G, Bordi C, de Bentzmann S. 2011. The PprA–PprB two-component system activates CupE, the first non-archetypal *Pseudomonas aeruginosa* chaperone–usher pathway system assembling fimbriae. Environ Microbiol 13:666–683.

118. Wang C, Chen W, Xia A, Zhang R, Huang Y, Yang S, Ni L, Jin F. 2019. Carbon Starvation Induces the Expression of PprB-Regulated Genes in *Pseudomonas aeruginosa*. Appl Environ Microbiol 85:e01705–19.

119. Barrow K, Kwon DH. 2009. Alterations in two-component regulatory systems of *phoPQ* and *pmrAB* are associated with polymyxin B resistance in clinical isolates of *Pseudomonas aeruginosa*. Antimicrob Agents Chemother 53:5150–5154.

120. Schurek KN, Sampaio JLM, Kiffer CRV, Sinto S, Mendes CMF, Hancock REW. 2009. Involvement of *pmrAB* and *phoPQ* in polymyxin B adaptation and inducible resistance in non-cystic fibrosis clinical isolates of *Pseudomonas aeruginosa*. Antimicrob Agents Chemother 53:4345–4351.

121. Fernández L, Jenssen H, Bains M, Wiegand I, Gooderham WJ, Hancock REW. 2012. The two-component system CprRS senses cationic peptides and triggers adaptive resistance in *Pseudomonas aeruginosa* independently of ParRS. Antimicrob Agents Chemother 56:6212–6222.

122. Muller C, Plésiat P, Jeannot K. 2011. A two-component regulatory system interconnects resistance to polymyxins, aminoglycosides, fluoroquinolones, and beta-lactams in *Pseudomonas aeruginosa*. Antimicrob Agents Chemother 55:1211–1221.

123. Bouvier J, Richaud C, Higgins W, Bögler O, Stragier P. 1992. Cloning, characterization, and expression of the *dapE* gene of *Escherichia coli*. J Bacteriol 174:5265–5271.

124. Fussenegger M, Facius D, Meier J, Meyer TF. 1996. A novel peptidoglycan-linked lipoprotein (ComL) that functions in natural transformation competence of *Neisseria gonorrhoeae*. Mol Microbiol 19:1095–1105.

125. Natale P, Brüser T, Driessen AJM. 2008. Sec- and Tat-mediated protein secretion across the bacterial cytoplasmic membrane—Distinct translocases and mechanisms. Biochim Biophys Acta Biomembr 1778:1735–1756.

126. Lan L, Murray TS, Kazmierczak BI, He C. 2010. *Pseudomonas aeruginosa* OspR is an oxidative stress sensing regulator that affects pigment production, antibiotic resistance and dissemination during infection. Mol Microbiol 75:76–91.

127. Kang Y, Nguyen DT, Son MS, Hoang TT. 2008. The *Pseudomonas aeruginosa* PsrA responds to long-chain fatty acid signals to regulate the *fadBA5*-beta-oxidation operon. Microbiology 154:1584– 1598.

128. Cox CD, Graham R. 1979. Isolation of an iron-binding compound from *Pseudomonas aeruginosa*. J Bacteriol 137:357–364.

129. Dietrich LEP, Teal TK, Price-Whelan A, Newman DK. 2008. Redox-active antibiotics control gene expression and community behavior in divergent bacteria. Science 321:1203–1206.

130. Fischer RS, Zhao G, Jensen RA. 1991. Cloning, sequencing, and expression of the P-protein gene (*pheA*) of *Pseudomonas stutzeri* in *Escherichia coli*: implications for evolutionary relationships in phenylalanine biosynthesis. J Gen Microbiol 137:1293–1301.

131. Goncharoff P, Nichols BP. 1984. Nucleotide sequence of Escherichia coli *pabB* indicates a common evolutionary origin of p-aminobenzoate synthetase and anthranilate synthetase. J Bacteriol 159:57– 62.

132. Bermingham A, Derrick JP. 2002. The folic acid biosynthesis pathway in bacteria: evaluation of potential for antibacterial drug discovery. Bioessays 24:637–648.

133. Reimmann C, Serino L, Beyeler M, Haa D. 1998. Dihydroaeruginoic acid synthetase and pyochelin synthetase, products of the *pchEF* genes, are induced by extracellular pyochelin in *Pseudomonas aeruginosa*. Microbiology (Reading) 144 (Pt 11):3135–3148.

134. Kurnasov O, Jablonski L, Polanuyer B, Dorrestein P, Begley T, Osterman A. 2003. Aerobic tryptophan degradation pathway in bacteria: novel kynurenine formamidase. FEMS Microbiol Lett 227:219–227.

135. Farrow JM, Pesci EC. 2007. Two Distinct Pathways Supply Anthranilate as a Precursor of the *Pseudomonas* Quinolone Signal. J Bacteriol 189:3425–3433.

136. Essar DW, Eberly L, Hadero A, Crawford IP. 1990. Identification and characterization of genes for a second anthranilate synthase in *Pseudomonas aeruginosa*: interchangeability of the two anthranilate synthases and evolutionary implications. J Bacteriol 172:884–900.

137. Crawford IP, Eberly L. 1986. Structure and regulation of the anthranilate synthase genes in *Pseudomonas aeruginosa*: I. Sequence of *trpG* encoding the glutamine amidotransferase subunit. Mol Biol Evol 3:436–448.

138. Lin J, Cheng J, Wang Y, Shen X. 2018. The *Pseudomonas* Quinolone Signal (PQS): Not Just for Quorum Sensing Anymore. Front Cell Infect Microbiol 8:230.

139. de Kievit TR, Iglewski BH. 2000. Bacterial Quorum Sensing in Pathogenic Relationships. Infect Immun 68:4839–4849.

140. Smith RS, Harris SG, Phipps R, Iglewski B. 2002. The *Pseudomonas aeruginosa* Quorum-Sensing Molecule N-(3-Oxododecanoyl)Homoserine Lactone Contributes to Virulence and Induces Inflammation In Vivo. J Bacteriol 184:1132–1139.

141. Cowell BA, Twining SS, Hobden JA, Kwong MSF, Fleiszig SMJ. 2003. Mutation of *lasA* and *lasB* reduces *Pseudomonas aeruginosa* invasion of epithelial cells. Microbiology 149:2291–2299.

142. Ortori CA, Dubern J-F, Chhabra SR, Cámara M, Hardie K, Williams P, Barrett DA. 2011. Simultaneous quantitative profiling of N-acyl-L-homoserine lactone and 2-alkyl-4(1H)-quinolone families of quorum-sensing signaling molecules using LC-MS/MS. Anal Bioanal Chem 399:839– 850.

143. Brewer LK, Jones JW, Blackwood CB, Barbier M, Oglesby-Sherrouse A, Kane MA. 2020. Development and bioanalytical method validation of an LC-MS/MS assay for simultaneous quantitation of 2-alkyl-4(1H)-quinolones for application in bacterial cell culture and lung tissue. Anal Bioanal Chem 412:1521–1534.

144. Yates EA, Philipp B, Buckley C, Atkinson S, Chhabra SR, Sockett RE, Goldner M, Dessaux Y, Cámara M, Smith H, Williams P. 2002. N-Acylhomoserine Lactones Undergo Lactonolysis in a pH-, Temperature-, and Acyl Chain Length-Dependent Manner during Growth of *Yersinia pseudotuberculosis* and *Pseudomonas aeruginosa*. Infect Immun 70:5635–5646.

145. Laville J, Blumer C, Schroetter CV, Gaia V, Défago G, Keel C, Haas D. 1998. Characterization of the *hcnABC* Gene Cluster Encoding Hydrogen Cyanide Synthase and Anaerobic Regulation by ANR in the Strictly Aerobic Biocontrol Agent *Pseudomonas fluorescens* CHA0. J Bacteriol 180:3187.

146. Folders J, Tommassen J, van Loon LC, Bitter W. 2000. Identification of a chitin-binding protein secreted by *Pseudomonas aeruginosa*. J Bacteriol 182:1257–1263.

147. Avichezer D, Gilboa-Garber N, Garber NC, Katcoff DJ. 1994. *Pseudomonas aeruginosa* PA-I lectin gene molecular analysis and expression in *Escherichia coli*. Biochim Biophys Acta 1218:11–20.

148. Winzer K, Falconer C, Garber NC, Diggle SP, Camara M, Williams P. 2000. The *Pseudomonas aeruginosa* Lectins PA-IL and PA-IIL Are Controlled by Quorum Sensing and by RpoS. J Bacteriol 182:6401–6411.

149. Wagner VE, Bushnell D, Passador L, Brooks AI, Iglewski BH. 2003. Microarray analysis of *Pseudomonas aeruginosa* quorum-sensing regulons: effects of growth phase and environment. J Bacteriol 185:2080–2095.

150. Ding F, Oinuma K-I, Smalley NE, Schaefer AL, Hamwy O, Greenberg EP, Dandekar AA. 2018. The *Pseudomonas aeruginosa* Orphan Quorum Sensing Signal Receptor QscR Regulates Global Quorum Sensing Gene Expression by Activating a Single Linked Operon. mBio 9:e01274–18.

151. Chugani SA, Whiteley M, Lee KM, D’Argenio D, Manoil C, Greenberg EP. 2001. QscR, a modulator of quorum-sensing signal synthesis and virulence in *Pseudomonas aeruginosa*. Proc Natl Acad Sci U S A 98:2752–2757.

152. Siehnel R, Traxler B, An DD, Parsek MR, Schaefer AL, Singh PK. 2010. A unique regulator controls the activation threshold of quorum-regulated genes in *Pseudomonas aeruginosa*. Proc Natl Acad Sci U S A 107:7916–7921.

153. Matsui H, Sano Y, Ishihara H, Shinomiya T. 1993. Regulation of pyocin genes in *Pseudomonas aeruginosa* by positive (*prtN*) and negative (*prtR*) regulatory genes. J Bacteriol 175:1257–1263.

154. Xiao G, Déziel E, He J, Lépine F, Lesic B, Castonguay M-H, Milot S, Tampakaki AP, Stachel SE, Rahme LG. 2006. MvfR, a key *Pseudomonas aeruginosa* pathogenicity LTTR-class regulatory protein, has dual ligands. Mol Microbiol 62:1689–1699.

155. Cao H, Krishnan G, Goumnerov B, Tsongalis J, Tompkins R, Rahme LG. 2001. A quorum sensing-associated virulence gene of *Pseudomonas aeruginosa* encodes a LysR-like transcription regulator with a unique self-regulatory mechanism. Proc Natl Acad Sci U S A 98:14613–14618.

156. Wade DS, Calfee MW, Rocha ER, Ling EA, Engstrom E, Coleman JP, Pesci EC. 2005. Regulation of *Pseudomonas* Quinolone Signal Synthesis in *Pseudomonas aeruginosa*. J Bacteriol 187:4372– 4380.

157. Knoten CA, Wells G, Coleman JP, Pesci EC. 2014. A Conserved Suppressor Mutation in a Tryptophan Auxotroph Results in Dysregulation of *Pseudomonas* Quinolone Signal Synthesis. J Bacteriol 196:2413–2422.

158. Palmer GC, Jorth PA, Whiteley M. 2013. The role of two *Pseudomonas aeruginosa* anthranilate synthases in tryptophan and quorum signal production. Microbiology (Reading) 159:959–969.

159. Wells IC. 1952. Antibiotic substances produced by *Pseudomonas aeruginosa*; syntheses of Pyo Ib, Pyo Ic, and Pyo III. J Biol Chem 196:331–340.

160. Saalim M, Villegas-Moreno J, Clark BR. 2020. Bacterial Alkyl-4-quinolones: Discovery, Structural Diversity and Biological Properties. Molecules 25:5689.

161. Wells G, Palethorpe S, Pesci EC. 2017. PsrA controls the synthesis of the *Pseudomonas aeruginosa* quinolone signal via repression of the FadE homolog, PA0506. PLoS ONE 12:e0189331.

162. Heinrichs JH, Gatlin LE, Kunsch C, Choi GH, Hanson MS. 1999. Identification and characterization of SirA, an iron-regulated protein from *Staphylococcus aureus*. J Bacteriol 181:1436–1443.

163. Cheung J, Beasley FC, Liu S, Lajoie GA, Heinrichs DE. 2009. Molecular characterization of staphyloferrin B biosynthesis in *Staphylococcus aureus*. Mol Microbiol 74:594–608.

164. Kuroda M, Hayashi H, Ohta T. 1999. Chromosome-determined zinc-responsible operon *czr* in *Staphylococcus aureus* strain 912. Microbiol Immunol 43:115–125.

165. Xiong A, Jayaswal RK. 1998. Molecular Characterization of a Chromosomal Determinant Conferring Resistance to Zinc and Cobalt Ions in *Staphylococcus aureus*. J Bacteriol 180:4024– 4029.

166. Balibar CJ, Shen X, McGuire D, Yu D, McKenney D, Tao J. 2010. *cwrA*, a gene that specifically responds to cell wall damage in *Staphylococcus aureus*. Microbiology 156:1372–1383.

167. Lin M-H, Li C-C, Shu J-C, Chu H-W, Liu C-C, Wu C-C. 2018. Exoproteome Profiling Reveals the Involvement of the Foldase PrsA in the Cell Surface Properties and Pathogenesis of *Staphylococcus aureus*. Proteomics 18:e1700195.

168. Lin M-H, Liu C-C, Lu C-W, Shu J-C. 2024. *Staphylococcus aureus* foldase PrsA contributes to the folding and secretion of protein A. BMC Microbiol 24:108.

169. Butala M, Zgur-Bertok D, Busby SJW. 2009. The bacterial LexA transcriptional repressor. Cell Mol Life Sci 66:82–93.

170. Smollett KL, Smith KM, Kahramanoglou C, Arnvig KB, Buxton RS, Davis EO. 2012. Global analysis of the regulon of the transcriptional repressor LexA, a key component of SOS response in *Mycobacterium tuberculosis*. J Biol Chem 287:22004–22014.

171. Agafonov DE, Spirin AS. 2004. The ribosome-associated inhibitor A reduces translation errors. Biochem Biophys Res Commun 320:354–358.

172. Smith EJ, Visai L, Kerrigan SW, Speziale P, Foster TJ. 2011. The Sbi Protein Is a Multifunctional Immune Evasion Factor of *Staphylococcus aureus*. Infect Immun 79:3801.

173. Bhakdi S, Tranum-Jensen J. 1991. Alpha-toxin of *Staphylococcus aureus*. Microbiol Rev 55:733.

174. Stapleton MR, Horsburgh MJ, Hayhurst EJ, Wright L, Jonsson I-M, Tarkowski A, Kokai-Kun JF, Mond JJ, Foster SJ. 2007. Characterization of IsaA and SceD, Two Putative Lytic Transglycosylases of *Staphylococcus aureus*. J Bacteriol 189:7316–7325.

175. Lang S, Livesley MA, Lambert PA, Littler WA, Elliott TS. 2000. Identification of a novel antigen from *Staphylococcus epidermidis*. FEMS Immunol Med Microbiol 29:213–220.

176. Cheung GYC, Rigby K, Wang R, Queck SY, Braughton KR, Whitney AR, Teintze M, DeLeo FR, Otto M. 2010. *Staphylococcus epidermidis* Strategies to Avoid Killing by Human Neutrophils. PLoS Pathogens 6:e1001133.

177. Cho H, Jeong D-W, Liu Q, Yeo W-S, Vogl T, Skaar EP, Chazin WJ, Bae T. 2015. Calprotectin Increases the Activity of the SaeRS Two Component System and Murine Mortality during *Staphylococcus aureus* Infections. PLoS Pathog 11.

178. Liu Q, Yeo W-S, Bae T. 2016. The SaeRS Two-Component System of *Staphylococcus aureus*. Genes (Basel) 7:81.

179. Bleul L, Francois P, Wolz C. 2021. Two-Component Systems of *S. aureus*: Signaling and Sensing Mechanisms. Genes 13:34.

180. Traber KE, Lee E, Benson S, Corrigan R, Cantera M, Shopsin B, Novick RP. 2008. *agr* function in clinical *Staphylococcus aureus* isolates. Microbiology (Reading) 154:2265.

181. Zapf RL, Wiemels RE, Keogh RA, Holzschu DL, Howell KM, Trzeciak E, Caillet AR, King KA, Selhorst SA, Naldrett MJ, Bose JL, Carroll RK. 2019. The Small RNA Teg41 Regulates Expression of the Alpha Phenol-Soluble Modulins and Is Required for Virulence in *Staphylococcus aureus*. mBio 10:e02484–18.

182. Briaud P, Zapf RL, Mayher AD, McReynolds AKG, Frey A, Sudnick EG, Wiemels RE, Keogh RA, Shaw LN, Bose JL, Carroll RK. 2022. The Small RNA Teg41 Is a Pleiotropic Regulator of Virulence in *Staphylococcus aureus*. Infect Immun 90:e0023622.

183. Romilly C, Lays C, Tomasini A, Caldelari I, Benito Y, Hammann P, Geissmann T, Boisset S, Romby P, Vandenesch F. 2014. A Non-Coding RNA Promotes Bacterial Persistence and Decreases Virulence by Regulating a Regulator in *Staphylococcus aureus*. PLoS Pathog 10:e1003979.

184. Menard G, Silard C, Suriray M, Rouillon A, Augagneur Y. 2022. Thirty Years of sRNA-Mediated Regulation in *Staphylococcus aureus*: From Initial Discoveries to In Vivo Biological Implications. Int J Mol Sci 23:7346.

185. Peschel A, Otto M, Jack RW, Kalbacher H, Jung G, Götz F. 1999. Inactivation of the *dlt* operon in *Staphylococcus aureus* confers sensitivity to defensins, protegrins, and other antimicrobial peptides. J Biol Chem 274:8405–8410.

186. Xue T, You Y, Hong D, Sun H, Sun B. 2011. The *Staphylococcus aureus* KdpDE Two-Component System Couples Extracellular K+ Sensing and Agr Signaling to Infection Programming. Infect Immun 79:2154–2167.

187. Michel A, Agerer F, Hauck CR, Herrmann M, Ullrich J, Hacker J, Ohlsen K. 2006. Global Regulatory Impact of ClpP Protease of *Staphylococcus aureus* on Regulons Involved in Virulence, Oxidative Stress Response, Autolysis, and DNA Repair. J Bacteriol 188:5783–5796.

188. Tuchscherr L, Bischoff M, Lattar SM, Noto Llana M, Pförtner H, Niemann S, Geraci J, Van de Vyver H, Fraunholz MJ, Cheung AL, Herrmann M, Völker U, Sordelli DO, Peters G, Löffler B. 2015. Sigma Factor SigB Is Crucial to Mediate *Staphylococcus aureus* Adaptation during Chronic Infections. PLoS Pathog 11:e1004870.

189. Pané-Farré J, Jonas B, Förstner K, Engelmann S, Hecker M. 2006. The σB regulon in *Staphylococcus aureus* and its regulation. Int J Med Microbiol 296:237–258.

190. Chien Y, Manna AC, Projan SJ, Cheung AL. 1999. SarA, a Global Regulator of Virulence Determinants in *Staphylococcus aureus*, Binds to a Conserved Motif Essential for *sar*-dependent Gene Regulation. J Biol Chem 274:37169–37176.

191. Morrison JM, Anderson KL, Beenken KE, Smeltzer MS, Dunman PM. 2012. The *Staphylococcal* Accessory Regulator, SarA, is an RNA-Binding Protein that Modulates the mRNA Turnover Properties of Late-Exponential and Stationary Phase *Staphylococcus aureus* Cells. Front Cell Infect Microbiol 2:26.

192. Zielinska AK, Beenken KE, Mrak LN, Spencer HJ, Post GR, Skinner RA, Tackett AJ, Horswill AR, Smeltzer MS. 2012. *sarA*-mediated repression of protease production plays a key role in the pathogenesis of *Staphylococcus aureus* USA300 isolates. Mol Microbiol 86:1183–1196.

193. Kim D, Paggi JM, Park C, Bennett C, Salzberg SL. 2019. Graph-based genome alignment and genotyping with HISAT2 and HISAT-genotype. Nat Biotechnol 37:907–915.

194. Lee DG, Urbach JM, Wu G, Liberati NT, Feinbaum RL, Miyata S, Diggins LT, He J, Saucier M, Déziel E, Friedman L, Li L, Grills G, Montgomery K, Kucherlapati R, Rahme LG, Ausubel FM. 2006. Genomic analysis reveals that *Pseudomonas aeruginosa* virulence is combinatorial. Genome Biol 7:R90.

195. Chung M, Adkins RS, Mattick JSA, Bradwell KR, Shetty AC, Sadzewicz L, Tallon LJ, Fraser CM, Rasko DA, Mahurkar A, Dunning Hotopp JC. 2021. FADU: a Quantification Tool for Prokaryotic Transcriptomic Analyses. mSystems 6:e00917–20.

196. Love MI, Huber W, Anders S. 2014. Moderated estimation of fold change and dispersion for RNA-seq data with DESeq2. Genome Biol 15:550.

197. Zhu A, Ibrahim JG, Love MI. 2019. Heavy-tailed prior distributions for sequence count data: removing the noise and preserving large differences. Bioinformatics 35:2084–2092.

198. Wu T, Hu E, Xu S, Chen M, Guo P, Dai Z, Feng T, Zhou L, Tang W, Zhan L, Fu X, Liu S, Bo X, Yu G. 2021. clusterProfiler 4.0: A universal enrichment tool for interpreting omics data. Innovation (Camb) 2:100141.

199. Yu G, Wang L-G, Han Y, He Q-Y. 2012. clusterProfiler: an R package for comparing biological themes among gene clusters. OMICS 16:284–287.

200. Li W, Godzik A. 2006. Cd-hit: a fast program for clustering and comparing large sets of protein or nucleotide sequences. Bioinformatics 22:1658–1659.

201. Winsor GL, Griffiths EJ, Lo R, Dhillon BK, Shay JA, Brinkman FSL. 2016. Enhanced annotations and features for comparing thousands of *Pseudomonas* genomes in the *Pseudomonas* genome database. Nucleic Acids Res 44:D646–D653.

202. Diep BA, Gill SR, Chang RF, Phan TH, Chen JH, Davidson MG, Lin F, Lin J, Carleton HA, Mongodin EF, Sensabaugh GF, Perdreau-Remington F. 2006. Complete genome sequence of USA300, an epidemic clone of community-acquired meticillin-resistant *Staphylococcus aureus*. Lancet 367:731–739.

